# Differential Avian Responses to Coffee Farming Lead to Community Homogenization in a Working Landscape in Jardín, Colombia

**DOI:** 10.1101/2025.02.21.639290

**Authors:** Fernando Cediel, Viviana Ruíz-Gutiérrez, Alejandra Echeverri, Juan. Luis Parra

## Abstract

In Colombia, coffee production has an ingrained cultural and economic importance. Coffee is cultivated in biodiverse areas using a variety of practices ranging from sun-exposed monocultures to mixed and shaded plantations. Management practices on this working landscape alter the composition of bird communities. We hypothesize that coffee agroecosystems homogenize bird communities by favoring the occupancy of non-forest dependent bird species and deterring forest dependent birds. We evaluate the effects of coffee plantations and associated metrics of vegetation structure on a subset of bird species in an agroforestry landscape in the Western Andes in Colombia and evaluate the hypothesis of community homogenization. We collected data in two consecutive years in 153 sites in coffee plantations and other habitats in Jardín, Colombia. In order to evaluate birds’ response to changes in the working landscape we made single-season occupancy models and used five covariates: coffee (binary covariate), foliage height diversity (FHD) and canopy height (CH) as proxies for coffee management practices, human footprint (LHFI), and elevation. Finally, we used generalized dissimilarity models (GDM) with the same covariates and geographical distance to evaluate the homogenization hypothesis among coffee sites. We identified 193 species (six endangered and five endemics) and modeled occupancy of 74 species. Coffee plantations significantly decreased the occupancy of 14 species while 13 were favored. In general, non-forest dependent species preferred lower FHD and lower CH values, and occurred at lower elevations with high LHFI. On the contrary, forest dependent species preferred higher FHD and CH, and were found at higher elevations. The GDM with best fit supported the hypothesis that coffee sites tend to be more similar than no-coffee sites, and that elevation and FHD explained most of the variation in dissimilarity. The working landscape in Jardín holds an important community of birds but coffee plantations are driving its homogenization. We anticipate that non-forest dependent birds will continue to colonize working landscapes, unless structural vegetation changes are promoted. Increasing FHD and CH is a potential strategy to improve forest dependent bird occupancy and reduce homogenization in Jardín.

## INTRODUCTION

Working landscapes cover approximately half of earth’s habitable surface (Ellis, et al., 2018). The distribution of crops on these landscapes is not random and often coincides with biodiversity hotspots. Therefore, it is important to understand how working landscapes can be designed to minimize harm to biodiversity (Kremen & Merelender, 2018). Coffee is among the working landscapes of economic importance in the Neotropics; 30% of the world’s coffee area is located in northern Latin America, from Mexico to Colombia (Vandermeer, 2003). Several studies have proposed coffee working landscapes as “beneficial” for conserving bird diversity since they can sustain similar bird communities as secondary forests (Bakermans, et al., 2012; Hernandez, et al., 2013; Frishkoff, et al., 2016), especially if these working landscapes are shaded coffee plantations embedded in a heterogeneous matrix (Perfecto & Armbrecht, 2003; Leyequién, et al., 2010; Bakermans, et al., 2012; Anderle, et al., 2023). Habitat heterogeneity and crop diversity support diverse biological communities (Isbell, et al., 2017; Guzman, et al., 2021), but some species do not maintain populations in the long term unless through colonization from nearby forest fragments (Frishkoff, et al., 2016; Hendershot, et al., 2023).

In South America, coffee plantations are in Andean slopes between 1000 and 2000 meters above sea level (masl) (Guhl, 2008), one of the most biodiverse and endangered regions in the world (Myers, et al., 2000; Kattan, et al., 2004; Kattan & Alvarez-López, 1996; Etter, et al., 2020). In Colombia, coffee plantations cover about 842.400 ha, of which 99% are technified coffee plantations (Fedecafé, 2022) meaning highly modified sun-exposed coffee farms (Moguel & Toledo, 1999) that have replaced native Andean forests (Etter, et al., 2006). Having sun-exposed coffee plantations is a relatively new practice (Borrero, 1986). In Colombia, most coffee was grown under native trees, called traditional rustic coffee (Moguel & Toledo, 1999). In the last few decades, intensification and technification of coffee plantations led to an abandonment of low intensity shaded coffee practices, including the replacement of coffee plants to a more resistant and productive coffee variety (Borrero, 1986; Guhl, 2008). This had negative impacts on biodiversity, and led to a higher pressure on adjacent remnant forests (Perfecto & Armbrecht, 2003; Karp, et al., 2018). Knowledge about biodiversity responses to heterogeneity within coffee agroecosystems is needed in order to reassess current efforts to balance economic and conservation goals.

Colombia is an ideal country to evaluate bird responses to coffee plantations due to its high avian diversity (Echeverry-Galvis, et al., 2022) and the cultural and economic importance of coffee (Guhl, 2008; Krishnan, 2017; UNESCO Heritage World, 2023). This relationship turns the process of cultivating coffee into a practice where a sense of place and cultural identity revolves around coffee. Colombian mountain ranges are ideal for coffee plantations (Guhl, 2008), and also for bird diversity (Kattan & Franco, 2004). Research has shown that coffee plantations with a tall and diverse canopy (e.g., traditional shade coffee systems), may at times support similar species richness relative to nearby native forests (Hernandez, et al., 2013). However, this apparent equivalence can be misleading, because most studies do not account for changes in community composition or increases in species richness that can happen as a result of species ability to exploit novel resources in disturbed landscapes (Komar, 2006; Karp, et al., 2018). These traditional coffee systems are lower yield relative to technified sun coffee systems. This conflict has motivated environmentally-minded coffee drinkers to support coffee brands that have been awarded certifications that account for more sustainable, regenerative, or Bird Friendly practices (Rueda & Lambin, 2013).

According to this, management decisions made by coffee-farm owners can have important consequences on biodiversity in and around coffee plantations (Mas & Dietsch, 2004; Philpott & Bichier, 2012; Valencia, et al., 2015). These decisions include, but are not limited to, the preservation of woodlands and waterways, fumigation and fertilization use and intensity and renovation of coffee plants and shade trees. In general, having more complex habitats with less intensity related to coffee production would help support the landscape-level persistence of the original bird community (Karp, et al., 2018). Past research has focused on understanding the role of coffee shade structure and its effects on avian community composition (Sánchez-Clavijo, Botero, & Espinosa, 2009; Jha, et al., 2014). However, less studies have evaluated how biological communities respond to different environmental policies, such as private-sector certification standards within agriculture, which can lead to differences in how farmers manage and steward the land.

Private-sector certifications, like Rainforest Alliance and Fair Trade, are a way to incentivize coffee farmers in reducing impacts on biodiversity while improving some social and economic aspects without decreasing coffee quality (Lentijo & Hostetler, 2013). Benefits to biodiversity include higher tree diversity, protecting waterways and land against erosion, among others, that convey ecosystem services like access to clean water, temperature regulation and wood provisioning (Lentijo & Hostetler, 2013; Perfecto, et al., 2005). Legislation can also promote the protection of biodiversity through policies that improve habitats for biodiversity (e.g., mainstreaming biodiversity into agriculture, (Echeverri, et al., 2023)). Although these certifications have been in the national scene for a long time, their impacts on biodiversity are not evident, since many of the implementations are not monitored and local environmental agencies don’t encourage communities to change their practices. Nonetheless, farms that follow the certifications’ standards tend to have richer and more diverse tree and bird communities (Mas & Dietsch, 2004; Philpott, et al., 2007). Certification standards are a potential tool to motivate farmers to protect biodiversity and improve their life quality without risking their income, but they need to be monitored to validate their use.

Changes in the composition of bird communities that arise in the process of working landscapes stem from changes in abundance, the appearance of new species, usually generalists, and the decrease of more specialized species (Drapeau, et al., 2000; Frishkoff, et al., 2019). It may take a long time for populations of species to be extirpated from working landscapes, especially if there are remnant forests nearby that help maintain the populations (Ferraz, et al., 2007). Thus, in a short time interval, the effect observed may be an increase in species richness, until forest dependent species are lost (Socolar, et al., 2016). In any case, the phenomenon observed is that in working landscapes the biodiversity tends to simplify and homogenize (Endenburg, et al., 2019; Velásquez-Trujillo, et al., 2021), especially where coffee plantations are managed intensively (i.e., sun-exposed coffee plantations). These effects are reduced where management practices are less intensive, for example, in working landscapes that follow certification standards. We predict that coffee plantations and other agroecosystems will favor habitat use by species that use open and less complex habitats (generalists). Simultaneously, although at a slower rate, agroecosystems should start to lose specialized species that are forest dependent (Socolar, et al., 2016; Karp, et al., 2018).

Here, we evaluate the effects of coffee plantations and management practices through indices of vegetation structure on avian habitat use and community composition in a Colombian working landscape. We used single-season occupancy models to evaluate the effects of important habitat characteristics (Foliage height diversity (FHD) and Canopy height (CH) as proxies for coffee management practices) on occupancy. We predicted heterogeneous bird responses to habitat variation. Forest dependent species should be negatively impacted by coffee plantations in relation to non-forest dependent species. We also predicted that bird composition among coffee sites should be more similar than among sites without coffee. Finally, based on bird responses in the agricultural landscape in Jardin, we offer guidelines on how to promote avian diversity through changes in management practices in coffee farms in the region.

## MATERIALS AND METHODS

### Study Area

This study was conducted in a working landscape in the vicinity of Jardin, a small town on the eastern slope of the western Andes (1750 masl) in the department of Antioquia, Colombia (Figure *1*).

**Figure 1.**
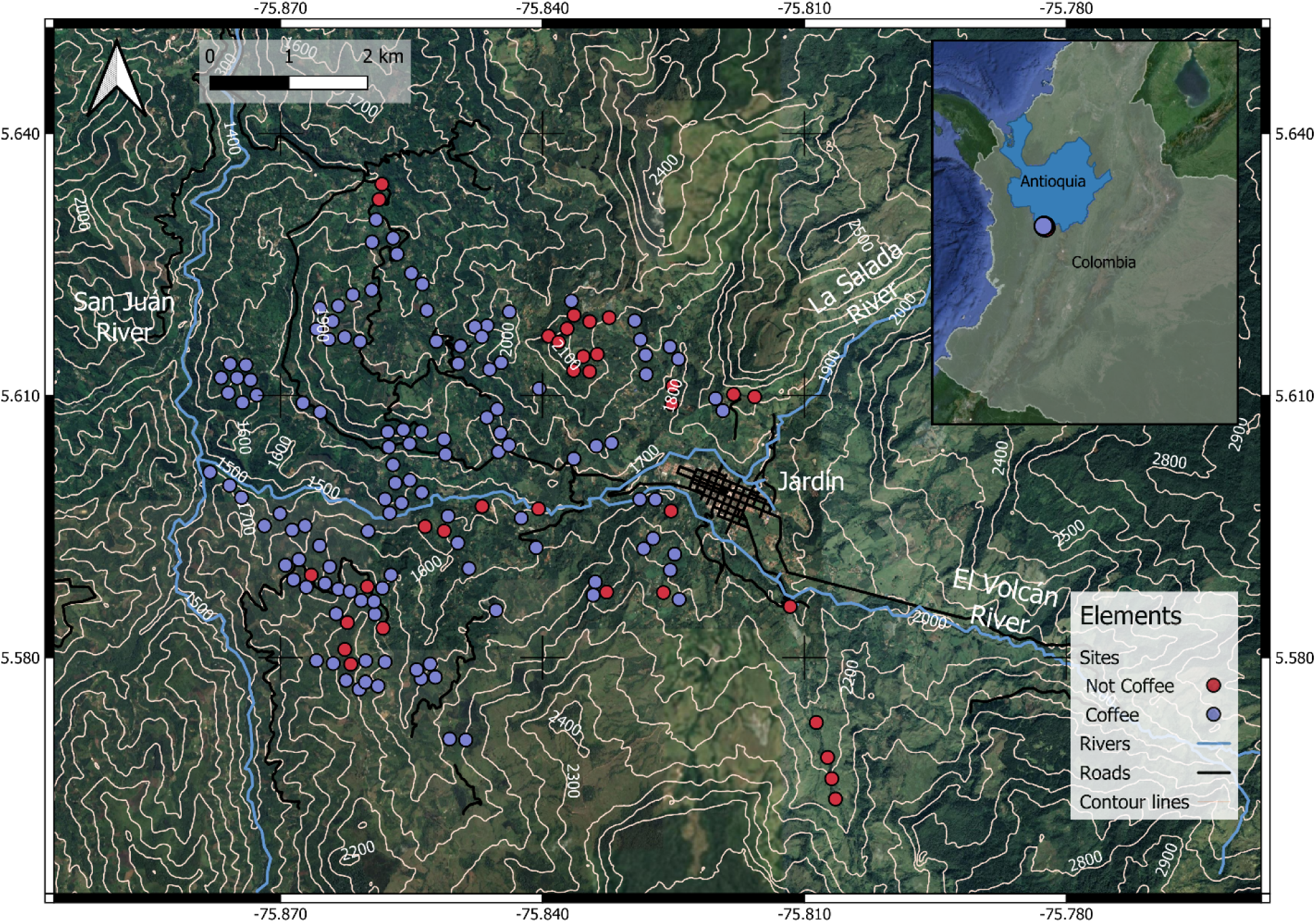
Study area indicating the point count localities (n=153) and the town of Jardín in the southwest corner of Antioquia. Red dots refer to point counts in non-coffee sites, such as forest patches (n=16), pastures (n=10), and other plantations (n=7 points), whereas purple dots refer to point counts on coffee farms (n=119).

This landscape has been highly modified and dominated by agroecological crops and pastures for several decades (Armenteras, Rodríguez, Retana, & Morales, 2011). Nonetheless, other plantations like sugarcane (*Saccharum* sp.), beans (*Phaseolus vulgaris*), avocado (*Persea americana*), various citrus trees and Eucalyptus (*Eucalyptus* sp.) are also common in the study area. Currently, montane forests in the study area are restricted to mountaintops, riversides and steep slopes, some of which are protected.

### Bird Sampling

To characterize bird communities across the landscape, we surveyed 153 sites distributed mostly in coffee farms and small roads. We followed the Proalas protocol (Ruiz-Gutiérrez, et al., 2020) designed to produce high-quality data for occupancy modeling in tropical habitats. Each site was surrounded by at least 100 m of the same habitat, was distanced at least 100 m from main roads and was separated by at least 200 m from other sampling sites to ensure independence. The majority of survey sites (n=119) were placed inside coffee plantations, as it is the predominant habitat in the landscape. The remaining sites were distributed across the least common habitats, like forest patches (16 points), pastures (10 points), and other plantations (7 points).

At least two experienced observers surveyed each site four times, during two years (2020 to 2021). Each year had two sampling seasons, one at the beginning of the year (January to March; dry season) and another one in the middle of the year (July to September; wet season). Surveys were performed during a four-hour period after dawn when birds’ activity is at its highest. We recorded every bird seen or heard and the distance at which they were detected (inside a 30 m or 100 m radius from the point count center). Each survey lasted between 10 and 15-min depending on the birds’ activity. All birds outside the 100 m radius, including gliding birds and highly mobile species (e.g., vultures and parrots) were not included because these would violate the assumption of independence among point count locations. To avoid temporal bias, we alternated the order of surveys every day. Survey data was entered via the eBird mobile application (Sullivan, et al., 2009).

### Data Analysis

To evaluate how birds responded to heterogeneity in the coffee landscape in Jardin, we used the observations from the 153 sites from the same season on 2020 and 2021 (each with 4 repetitions during the dry season) and employed single species single-season occupancy models (MacKenzie, et al., 2002). We concentrated our analyses on this subset of the data since it was the most stable in terms of observer experience (i.e., observers had sufficient experience with the region’s birds to know their vocalizations and visual identification cues) and climate (i.e., no precipitation anomalies due to ENSO related events). Since our detection histories were spread over two years and occupancy models assume that occupancy state is constant during a season, we duplicated the number of sites and assigned the detection histories from the second year to the second set of sites increasing sample size to better estimate occupancy relationships. In other words, we assume that data from the second year are independent from the first year and allow the model to estimate occupancy with an increased sample size. To promote model convergence and avoid species with extremely low detection, we only included bird species that were recorded in at least 10 sites during the two seasons. Each species was assigned a forest-dependency category based on the Birdlife International classification (Birdlife International, 2023).

We used the following detection covariates that we recorded at every survey visit: observation time, landscape noise, and season. Observation time (hours and minutes) is the time in a decimal format at the start of each survey (hour plus minutes/60, e.g., 6.583 to indicate 06:35). Landscape noise was measured in decibels with an application for smartphones (Sound Meter at Google play store) for at least 5 minutes during the survey and the value was written on the eBird’s list comments section. Season was a binary covariate denoting the year when the survey was made (0 = dry 2020, 1 = dry 2021). Occupancy covariates included: Canopy Height (CH), which represents the mean height of the tree canopy, Foliage Height Diversity (FHD) which captures vegetation’s vertical complexity and is related to habitat quality and bird diversity (Halma, 1975). CH and FHD were derived from the GEDI lidar spaceborne sensor at a 30 m spatial resolution for the year 2019 (Hakkenberg, et al., 2023) and processed for Colombia by Fagua et al. (2021). Legacy-adjusted human footprint index (LHFI) is a metric that represents all human interventions (e.g., infrastructure, mining projects, population density) for the year 2015 and was obtained from Correa Ayram et al. (2020) at 300 m spatial resolution, Elevation (EL) was obtained from the Digital elevation Model at http://srtm.csi.cgiar.org developed by Reuter et al. (2007) at 90 m resolution (Figure S1). The last occupancy covariate indicated whether the site was in a coffee plantation (1=Coffee) or not (0). All covariates, except coffee, were assigned to each point count locality from raster archives using a 100 or 300 m buffer depending on the raster resolution.

All analyses were made in R statistical software (R Core Team, 2022). We checked for collinearity among covariates through a correlation analysis and found that none of the variables were significantly associated with each other (Pearson’s r < 0.8), so we used them all in the models (Figure S2). We standardized the covariates to a mean of 0 and variance of one prior to model fitting and we used the package “unmarked” (Fiske & Chandler, 2011) to fit the occupancy models. For every species, we first identified the best detection model among five candidates (Table *1*) while holding occupancy constant (MacKenzie, et al., 2017; Morin, et al., 2020).

**Table 1.** Detection models used in increasing order of complexity (number of parameters). Model notation includes the covariates used within parentheses for occupancy (Ψ) and detection (ρ). Points within parentheses indicate models without covariates. Covariates include the noise measured in decibels (Decibels), the time at the start of the point count (Time), and the year when the point count was made (Season). Full model included all three covariates.

| Model | Covariates | Model Name | # Parameters |
| --- | --- | --- | --- |
| $\psi(.)\rho(.)$ | Null | Null | 2 |
| $\psi(.)\rho(\text{decibels})$ | Decibels | Noise | 3 |
| $\psi(.)\rho(\text{time})$ | Time | Time | 3 |
| $\psi(.)\rho(\text{season})$ | Season | Season | 3 |
| $\psi(.)\rho(\text{decibels.time.season})$ | Full | Full | 5 |

After identifying the best detection model, we evaluated a set of 15 occupancy models (Table *2*) using the best detection model. We excluded interactions between covariates to simplify interpretation and avoid overparameterization (Ortega-Álvarez, et al., 2020). We excluded models that did not converge, or that showed standard error estimates higher than 1 for any of the coefficients. Then, we ranked models using Akaike Information Criterion corrected by sample size (AICc) (Burnham & Anderson, 2004) with the “MuMIn’’ package (Barton, 2023). Finally, we did model-averaging with the resulting models to better account for uncertainty among models for all species. We averaged the occupancy models using the “modavg” function in the “AICcmodavg’’ package (Mazerolle., 2023).

**Table 2.** Occupancy models in increasing order of complexity by the number of parameters used for each species. Model notation includes the covariates used within parentheses for occupancy (Ψ) and detection (ρ). Points within parentheses indicate models without covariates. Covariates include the height of the canopy (CH), foliage height diversity (FHD), elevation (EL), human footprint (LHFI) and a categorical covariate that states if a point count was in a coffee plantation (Coffee) or not.

| Model | Covariates | Model Name | # Parameters |
| --- | --- | --- | --- |
| $\psi(.)\rho(\text{detection covariates})$ | None | null | 2 |
| $\psi(\text{CanHei})\rho(\text{detection covariates})$ | Canopy height | height | 3 |
| $\psi(\text{FHD})\rho(\text{detection covariates})$ | FHD | fhd | 3 |
| $\psi(\text{Elev})\rho(\text{detection covariates})$ | Elevation | elev | 3 |
| $\psi(\text{LHFI})\rho(\text{detection covariates})$ | Legacy-adjusted Human Footprint Index | lhfi | 3 |
| $\psi(\text{Coffee})\rho(\text{detection covariates})$ | Coffee categories | cat | 3 |
| $\psi(\text{LHFI})\rho(\text{detection covariates})$ | Legacy-adjusted Human Footprint Index | height.fhd | 5 |
| $\psi(\text{FHD.Elev})\rho(\text{detection covariates})$ | Foliage height diversity + Elevation | fhd.elev | 5 |
| $\psi(\text{CanHei.Elev})\rho(\text{detection covariates})$ | Canopy height + Elevation | height.elev | 5 |
| $\psi(\text{CanHei.LHFI})\rho(\text{detection covariates})$ | Canopy height + Legacy-adjusted Human Footprint Index | height.lhfi | 5 |
| $\psi(\text{CanHei.Coffee})\rho(\text{detection covariates})$ | Canopy height + Coffee categories | height.cat | 5 |
| $\psi(\text{FHD.Coffee})\rho(\text{detection covariates})$ | Foliage height diversity + Coffee categories | fhd.cat | 5 |
| $\psi(\text{Elev.Coffee})\rho(\text{detection covariates})$ | Elevation + Coffee categories | elev.cat | 5 |
| $\psi(\text{LHFI.Coffee})\rho(\text{detection covariates})$ | Legacy-adjusted Human Footprint Index + Coffee categories | lhfi.cat | 5 |
| $\psi(\text{CanHei.FHD.Elev.LHFI.Coffee})\rho(\text{detection covariates})$ | Canopy height + Foliage height diversity + Elevation + Legacy-adjusted Human Footprint Index + Coffee categories | full | 7 |

To evaluate the biotic homogenization hypothesis, we used Generalized Dissimilarity Modeling (GDM; (Fitzpatrick, et al., 2021)), a regression technique that predicts the rate of species turnover (i.e., beta diversity) as a function of several distances including geographic and environmental distances and uses monotonic *I*-splines to show the effects of each covariate on beta diversity through their height and slope (Ferrier, et al., 2007). We quantified betadiversity using Bray Curtis distance based on presence/absence data and used euclidean geographic distance (geo dist), and environmental distances based on each of the following covariates: canopy height (CH), foliage height diversity (FHD), legacy-adjusted human footprint index (LHFI), elevation (EL), and whether the site was in a coffee plantation (Coffee). We evaluated the relative importance among covariates through the percent decrease in deviance explained between the full model and a model fitted with that predictor permuted, and their significance using matrix permutation with the function “gdm.varImp’’ from the R package “gdm” (Fitzpatrick, Mokany, Manion, Nieto-Lugilde, & Ferrier, 2021).

To evaluate the hypothesis that coffee plantations promote biotic homogenization, (i.e., decreased beta diversity), we tested the following alternatives, assigning categorical distances to the three possible comparisons between sites (coffee-coffee, not coffee - not coffee, and coffee - not coffee). First, as a null model, we excluded the coffee covariate, and evaluated two alternatives: (1) coffee has the same homogenizing effect as other categories: we assigned site comparisons in the same category as 0 distance, and comparisons between sites in different categories a distance of 1, and (2) coffee plantations homogenize bird diversity: smallest difference (0) was assigned to comparisons between coffee sites, and a higher difference was assigned to comparisons between sites without coffee (1), and the largest difference (2) assigned to comparisons between coffee and not coffee sites. Finally, based on the deviance explained, we identified the alternative that best suited our observations. We made GDMs with the modeled occupancies (corrected for imperfect detection) of the 74 species with sufficient detections. Since this approach uses model predictions that are based on environmental covariates, we use results of the GDM to evaluate their overall contribution to betadiversity when accounting for imperfect detection. We also made GDMs based on the complete bird community using raw observations (see supplementary material). As a final contribution to better understand which species were driving betadiversity patterns, we made a similarity percentage analysis with the “simper” function of the package “vegan” (Oksanen, et al., 2022) in R.

## RESULTS

### General Results

We collected 7610 detections representing 193 species during the two seasons between 2020 and 2021. Among these species, 50 are of interest for conservation (Table S1). Five species are endemic to Colombia and 14 near-endemic. Six species are in red list categories: two near-threatened (NT), three vulnerable (VU) and one endangered (EN). We also detected 14 boreal migratory species (Echeverry-Galvis, et al., 2022). Of the 193 species, we assigned 125 that had medium or high forest dependency to the forest dependent category, while the rest (68) were classified as non-forest dependent. For the occupancy analyses, we selected 86 species that were detected at ten or more sites. In this group, ten species are of interest for conservation, four are endemic and six near-endemic. 46 species are in the forest dependent category, and 40 are classified as non-forest dependent (Table S1).

### Occupancy Models

We were able to successfully model the selected 74 species and found a gradient of responses across species (Figure 2). Non-forest dependent species, like the Colombian Chachalaca (*Ortalis columbiana*) preferred lower elevations (mean occupancy coefficient −0.575 SE 0.370) and a higher LHFI (mean occupancy coefficient 0.282 SE 0.314). On the other hand, forest dependent species like the Scaled Antpitta (*Grallaria guatimalensis*) and the Moustached Puffbird (*Malacoptila mystacalis*) showed a preference for sites with lower LHFI (mean occupancy coefficient −0.108 SE 0.358) and higher elevations (mean occupancy coefficient 0.077 SE 0.377) Figure S3 and Figure S4).

**Figure 2.**
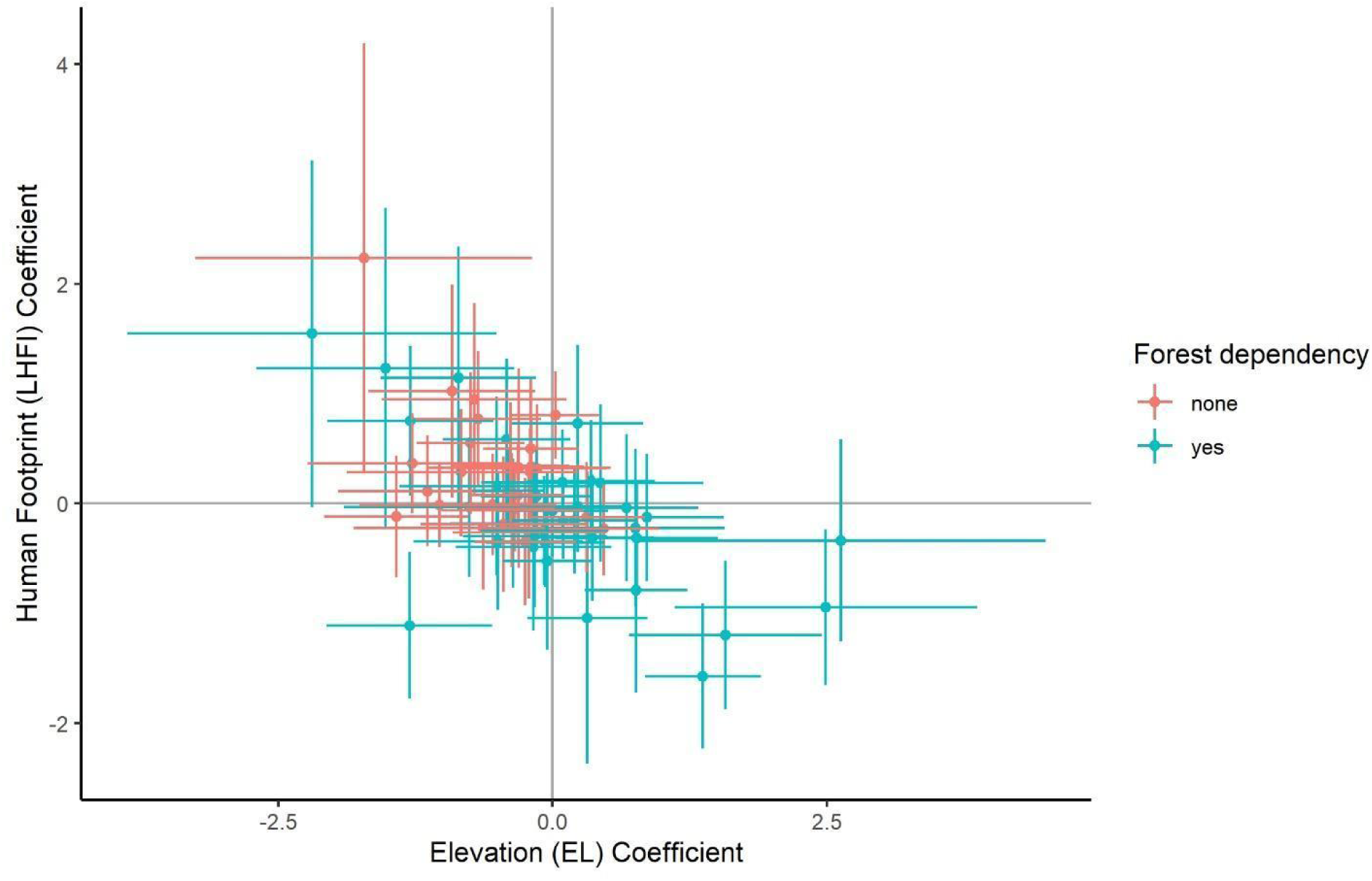
Occupancy coefficients for all 74 species modeled with its corresponding confidence 95% intervals for elevation and human footprint. Colors represent forest and non-forest-dependent species. Coefficients are in logit scale.

We found 24 species in which coffee appeared as a covariate among plausible models, including 12 species with regression coefficients for the effect of coffee whose 95% confidence interval did not overlap 0 (Figure 3). Eight of these species decreased their occupancy in coffee plantations, among these, the endemic Parker’s Antbird (*Cercomacroides parkeri*). Two near-endemic species, Whiskered Wren (*Pheugopedius mystacalis*) and Black-winged Saltator (*Saltator atripennis*), had negative but non-significant negative responses. Six of the 7 species that were negatively affected by coffee plantations are forest dependent species and their occupancy increased with higher FHD (Figure 4). On the other hand, 3 species showed a significant positive response to coffee plantations, all of which are non-forest dependent and who have a positive response in occupancy with lower FHD.

**Figure 3.**
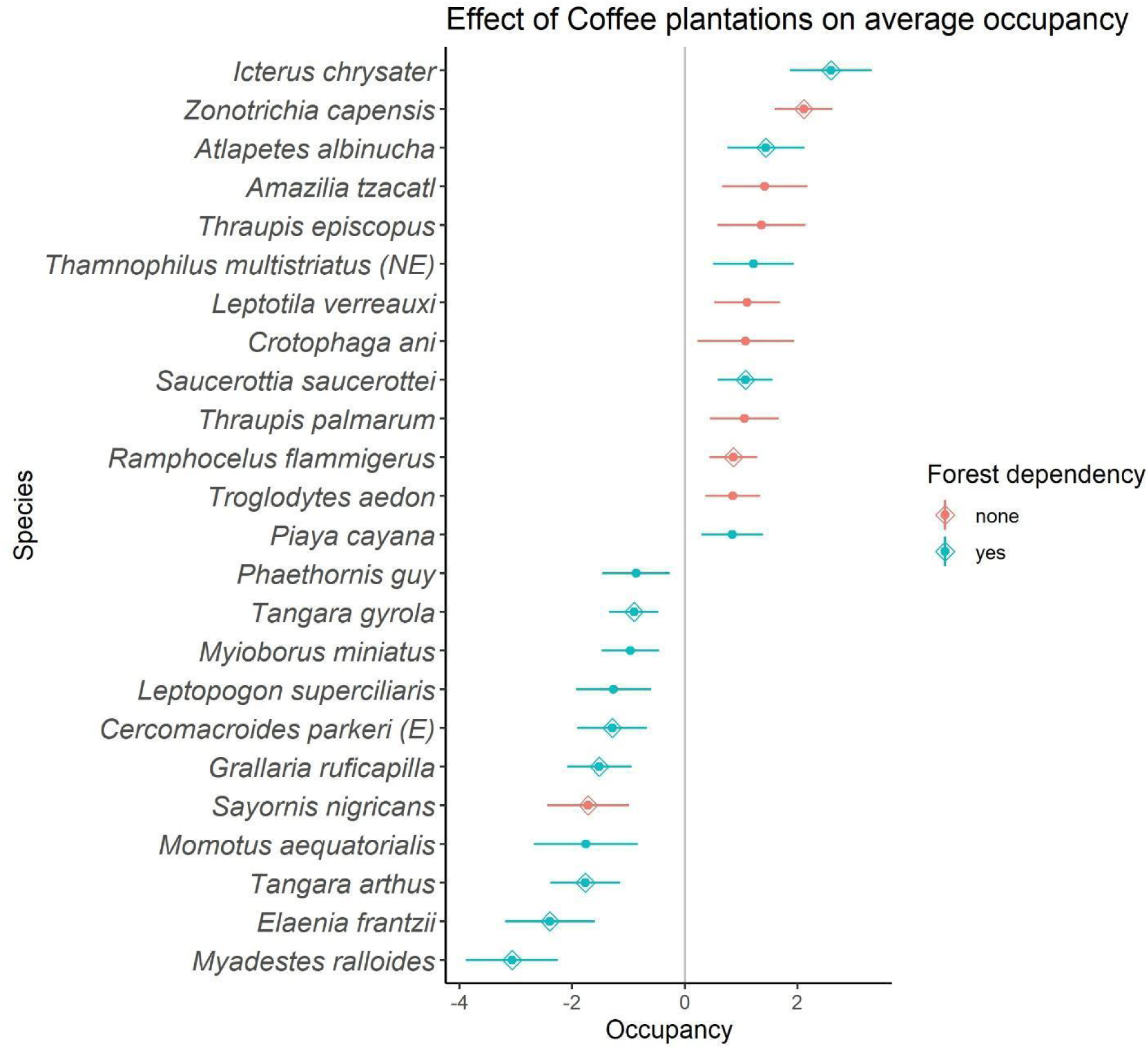
Species impacted by coffee plantations. This set of 24 species includes those whose standard errors for the model averaged coefficients for coffee did not overlap with 0. Species whose 95% confidence interval for the coffee coefficient did not overlap 0 are marked with a diamond. Colors indicate forest dependency as classified by Birdlife International. Coefficients are in logit scale.

**Figure 4.**
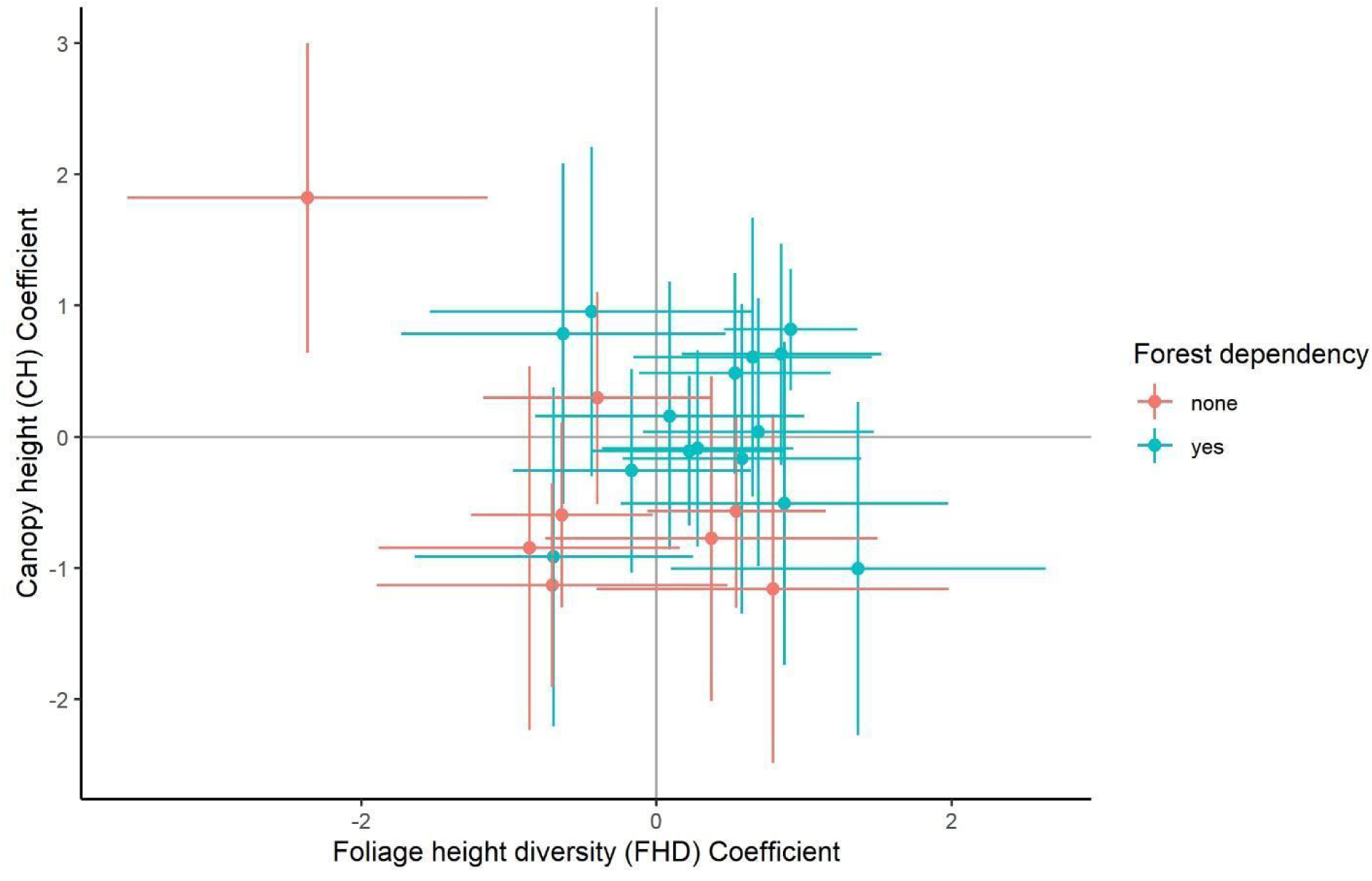
Occupancy coefficients and their corresponding 95% confidence intervals of foliage height diversity and canopy height for 24 species whose occupancy is affected by coffee plantations (see Figure 3). Colors represent forest dependent species as classified by Birdlife international. Coefficients are in logit scale.

Of these 24 species (Figure 4), we checked the effect of both habitat covariates (CH and FHD) on occupancy and found that most non-forest dependent species (62.5%) prefer habitats with lower-than-average FHD (left). On the contrary, most forest dependent species (53.3%) prefer sites with a higher-than-average FHD, but some with high or low canopies.

### General Dissimilarity Model (GDM)

The GDM based on occupancy predictions of the 74 modeled species had an explained deviance of ∼93%. The GDM identified Elevation (relative predictor importance = 35.4), followed by FHD (10.1), CH (3.4) and LHFI (5.5), as the covariates that predicted the most variation in compositional dissimilarity. Whether or not the site was on a coffee plantation had a relative low effect on ecological dissimilarity (Predictor importance = 2.3). Geographic distance did not add any explanatory power to the GDM. The GDM that used raw detections from all species had an explained deviance of 33%, and Elevation was the most informative variable (26.7), followed by FHD (11.4), coffee categories (6.1), Geographic distance (4.5) and lastly LHFI (0.8), explained variation in compositional dissimilarity, whereas CH had no contribution (Figure S5 and Figure S6).

In relation to the effects of coffee as a driver of biotic homogenization, we found that excluding the coffee covariate in the analysis decreased the deviance explained in both versions of the GDM (Table *3*). The alternative that supported the biotic homogenization hypothesis (coffee-coffee site comparisons most similar) increased the deviance explained in the GDM with the complete community data (Figure S5), but not relative to the equal effects hypothesis in the GDM that used predicted occupancies for a subset of the species (Figure 5).

**Figure 5.**
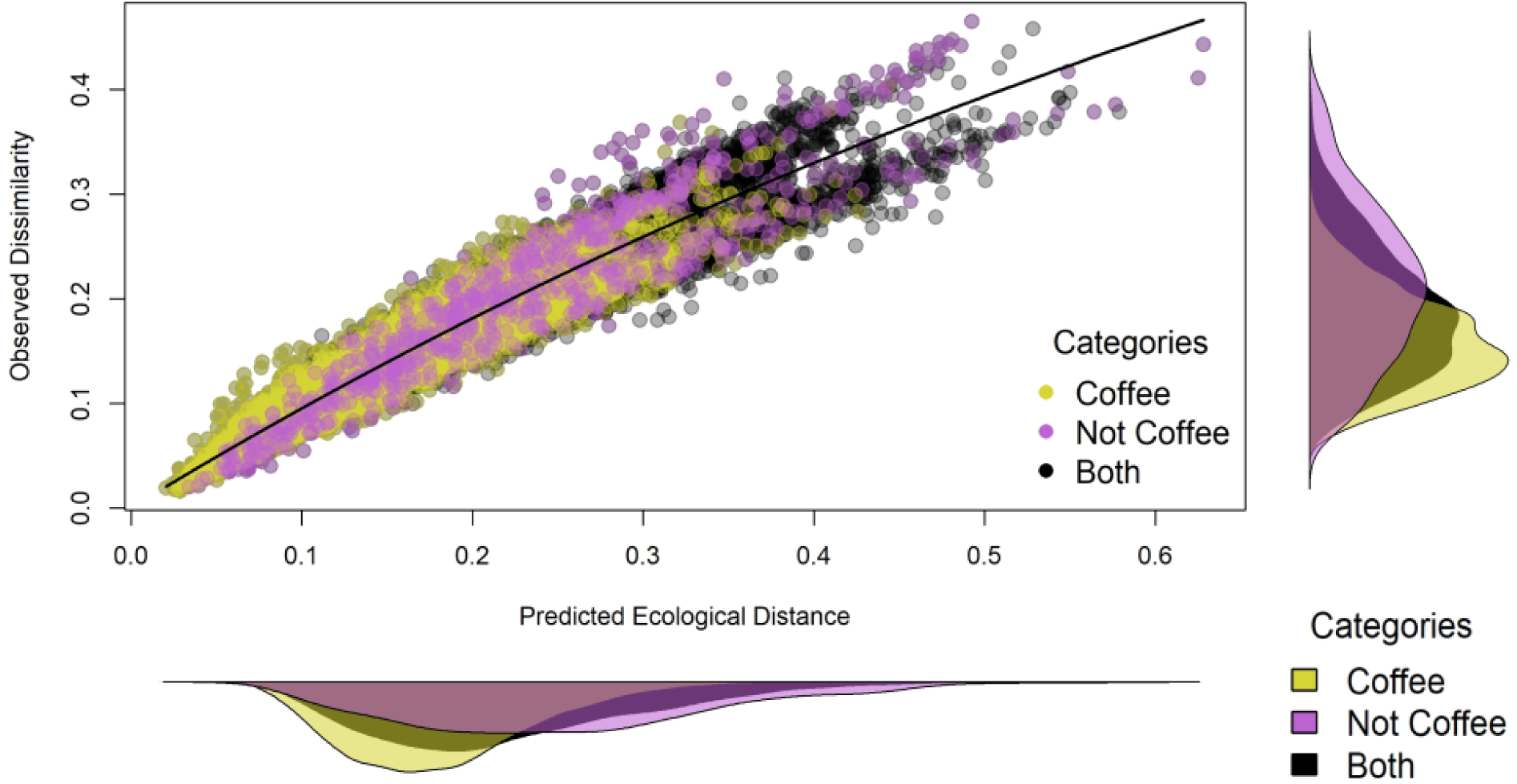
Observed compositional dissimilarities (based on Bray-Curtis distance among communities using modeled occupancies for 74 species) as a function of predicted ecological distances among sites using all covariates. Each point represents a site-pair comparison and is colored depending on the nature of the comparisons (coffee *–* coffee, non-coffee *–* non-coffee, and coffee *–* non-coffee). Side graphs show densities of each axis.

**Table 3.** GDM explained deviance for the three alternative hypotheses about the role of coffee plantations for betadiversity on the two approches: i. Null model does not include coffee as a covariate in the GDM, ii. coffee – coffee comparisons are assigned the same ecological difference as non-coffee – non-coffee, iii. coffee – coffee comparisons are assigned the highest ecological difference (biotic homogenization hypothesis).

| GDM approach | Null model (excluding coffee) | Equal effects | Biotic homogenization (due to coffee) |
| --- | --- | --- | --- |
| Complete community | 29,347 | 31,256 | 33,246 |
| Modeled species | 91,091 | 93,322 | 93,295 |

The shape of the elevation spline function in the GDM indicates that predicted ecological distance increases steadily with elevation, whereas predicted ecological distance due to FHD is higher at higher values of FHD (>2.8) than at lower values of FHD (<2.6). Intermediate values of CH (18-22 m) and LHFI (55-60) resulted in the largest changes in ecological distance (Figure 5). Comparisons between coffee and not-coffee sites had the largest contribution to ecological distance, and geographic distance had no effect (Figure 6). The shapes and relative importance of the spline functions were considerably different in the GDM that used all species raw detections (Figure S6). The relative importance changes on the two approximations, geographic distance changes on the two analyses.

**Figure 6.**
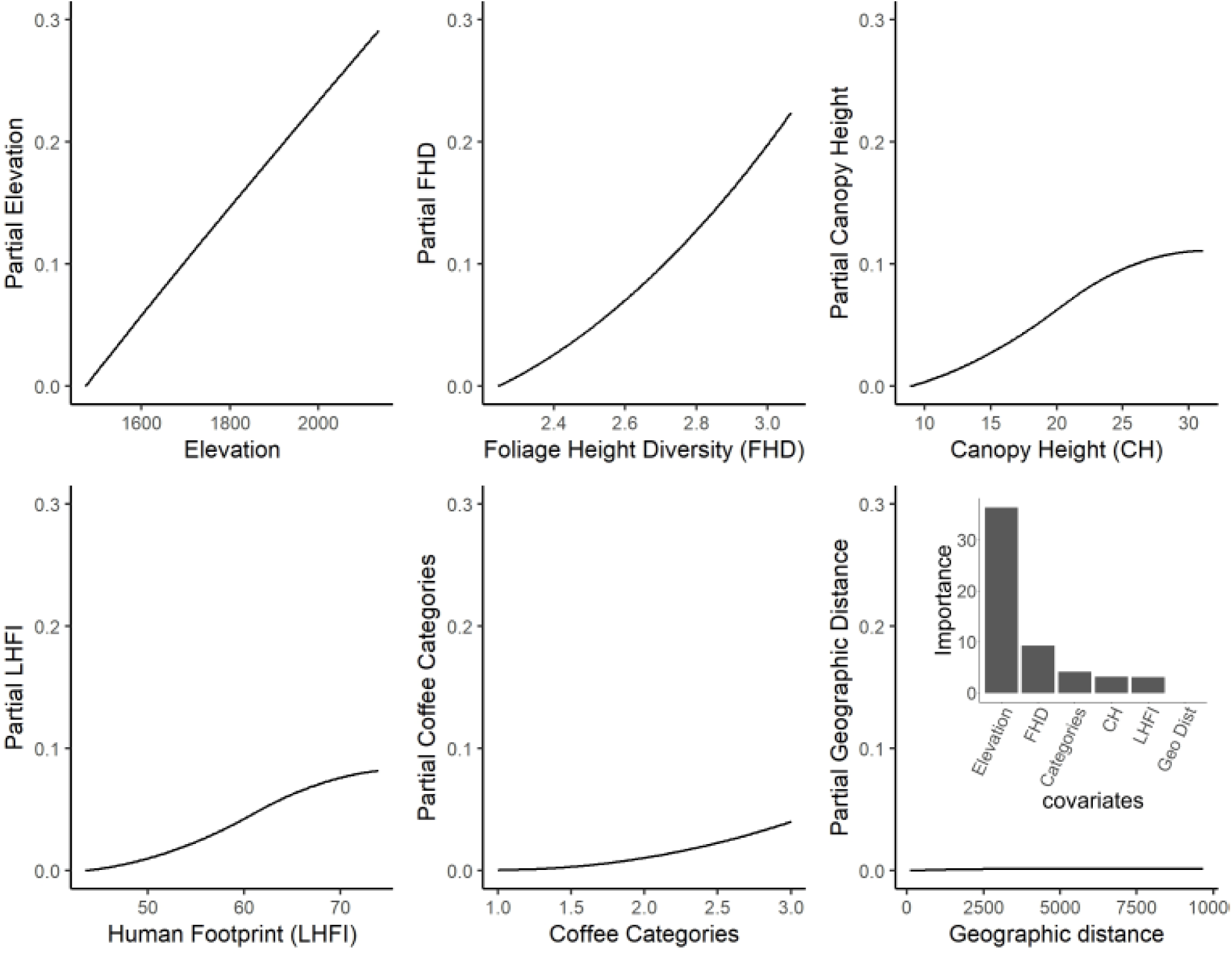
I-splines for each covariate used in the GDM analysis. The I-splines show the relationship between each variable and the predicted ecological distance. In this case, geographic distance (Geo Dist) was the least important predictor variable. The barplot on the lower right is the result of the relative importance of the covariates used.

The species that most contributed to dissimilarity between coffee and non-coffee sites when using modeled occupancy (Table S2) was the Golden Tanager (*Tangara arthus*), a forest dependent species whose average abundance was higher in non-coffee than in coffee sites. The next species was the Bananaquit (*Coereba flaveola*), a non-forest species whose average abundance at coffee sites was higher than in non-coffee sites. The overall contribution of forest dependent species to the dissimilarity between coffee and non-coffee sites was 0.48 (using modeled occupancy)(Table S2 and Table S3).

## DISCUSSION

Our results indicate that coffee plantations in Jardín have a significant and differential effect on bird occupancy. At the community level, coffee plantations seem to homogenize bird composition by filtering out forest dependent species and promoting the establishment of non-forest dependent species. Our key findings support conclusions from other studies about bird diversity in agroecosystems (Calvo & Blake, 1998; Tejeda-Cruz & Sutherland, 2004; Velásquez-Trujillo, et al., 2021) and offer a potential simple strategy to promote the occupancy of forest dependent birds including endemics, threatened, and migratory species: an increase in foliage height diversity (FHD). Coffee plantations in Jardin have a negative effect on most forest dependent bird species, and a positive effect on non-forest dependent bird species (Figure 3, Figure *4* & Figure S 4), in line with results from other studies (Valente, et al., 2022). We also found that simplification of vegetation structure, which is expected to occur in homogeneous unshaded monoculture plantations (e.g., sun-exposed coffee), favors non-forest dependent birds, while a more complex vegetation structure (rustic or polyculture coffee) promotes occupancy of forest dependent birds (Calvo & Blake, 1998). The working landscape studied in Jardin, like other working landscapes in the Andean region, is characterized by a mixture of coffee plantation practices including sun exposed, mixed plantations, shaded, and variations of these (Moguel & Toledo, 1999). Despite this landscape heterogeneity, our results indicate that bird species composition is more similar between coffee plantations relative to sites without coffee.

Coffee plantations with diverse shade and close to natural habitats have shown to be suitable for forest dependent birds (Borrero, 1986; Jha, et al., 2014). Our results show that the working landscape in Jardín holds an important bird community which includes endemics and threatened forest dependent species (Table S1). This coincides with results from other coffee agroecosystems in Colombia (Valente, et al., 2022) and other countries (Calvo & Blake, 1998; Kammerichs-Berke, et al., 2022). Nevertheless, it is important to emphasize that not all birds respond in a similar manner to changes in land use (Contreras, et al., 2022; Valente, et al., 2022). Our results show that the occupancy of eight non-forest dependent species increases at coffee plantations (Figure S4). Six of those eight species also increase their occupancy at sites with a higher human footprint and lower altitudes, and five of those species also prefer sites with lower foliage height diversity (e.g., shaded monoculture). Most of these species prefer warmer sites and some are apparently expanding their distribution in Colombia aided by deforestation and climate change (Londoño Gómez & Pulgarín-Restrepo, 2024; Avendaño, et al., 2013). On the contrary, occupancy of forest dependent species including understory and canopy species declined in coffee plantations with low foliage height diversity (i.e., sun-exposed coffee). These opposing trends imply that generalist species - for example, the White-tipped Dove (*Leptotila verreauxi),* Palm Tanager (*Thraupis palmarum*), and Smooth-billed Ani (*Crotophaga ani*) - may be colonizing coffee plantations from lower elevations, (Connell, 1977; Valente, et al., 2022), while more forest-specialized species - for example, the Andean Solitaire (*Myadestes ralloides*), Parkeŕs Antbird (*Cercomacroides parkeri*) and Chestnut-crowned Antpitta (*Grallaria ruficapilla*) - might be vanishing because of habitat reduction and fragmentation as well as climate change (Malavasi, et al., 2009; Velásquez-Trujillo, et al., 2021). The shifting balance between these two mechanisms may be responsible for the diversity of results regarding avifauna in agroecosystems.

Our results suggest that agroecosystems in Jardín, despite maintaining an important bird community, are driving its homogenization. In general, it is expected that declining heterogeneity in the landscape decreases betadiversity (Benton, et al., 2003; Velásquez-Trujillo, et al., 2021) and coffee monocultures (shaded or unshaded) are mostly decreasing landscape heterogeneity in Latin America (Moguel & Toledo, 1999). The Andes are characterized by high levels of betadiversity across subregions (Kattan et al., 2004), a pattern that endorses the conservation of small patches spread out rather than a single large patch (Kattan & Alvarez-López, 1996). Results of the GDM show that coffee plantations are not the main contributor to the turnover in bird composition across the landscape. The dissimilarity between two coffee sites was on average lower than the dissimilarity between a coffee and a non-coffee site (Figure 5 & Figure S5). On the other hand, the highest dissimilarity, on average, was among non-coffee sites. This shows that maintaining heterogeneous coffee landscapes that have other habitats besides coffee will help prevent homogenization (Karp, et al., 2018; Valente, et al., 2022). Our results also show that coffee plantations in Jardín sustain low levels of betadiversity. This is worrying since the tendency in this region is to expand intense coffee plantations (unshaded monoculture).

The rate of decay of forest dependent species in agroecosystems like coffee plantations, might be low, especially if the landscape continues to hold scattered forest fragments. Thus, at certain time-frames and spatial extents, a complex agroecosystem landscape may hold more species than a forest (Hernandez, et al., 2013; Estrada-Carmona, et al., 2019), but this should not be taken as evidence that agroecosystems benefit biodiversity. If landscape heterogeneity continues to decrease, this might pave the way for other non-forest dependent, generalistic species to arrive and eventually outcompete former species (Socolar, et al., 2016). One potential solution arising from our results is to increase FHD and CH in coffee plantations, which would favor the occupancy of some forest-dependent species. Another solution is to increase landscape heterogeneity, for example, through strategies like land-sparing and land-sharing (Kremen, 2015).

According to the *I-*splines from the best GDM model, community turnover in Jardín is driven by variation in elevation (EL), foliage height diversity (FHD), and canopy height (CH); the former two had the strongest effects on betadiversity. Bird community turnover occurred continuously across the elevation gradient. This result is consistent with several studies in tropical mountains at intermediate elevations (Jankowski, et al., 2013; Presley, et al., 2012). FHD had a stronger effect on ecological dissimilarity at higher values (> 2.8), suggesting that sites with the highest vertical foliage diversity had the most distinct composition (Moorhead *et al*. 2010). In general, our results indicate that having a more complex vertical structure is associated with greater bird diversity (Cruz-Angón & Greenberg, 2005). Canopy height explained bird turnover but its effect fainted at about 25 m, where taller canopies had no further effect on turnover. This could be because the sites with the tallest canopies, refer to small forests sites or coffee plantations surrounded by tall trees (e.g., Eucalyptus or Pines) that are used to mark the roads or the properties, which do not hold such an important bird community (Marsden, et al., 2001). This configuration (shaded monoculture) can be thought of as a monoculture above another monoculture, lacking structural complexity, although it could have some positive effects like carbon sequestration (Ehrenbergerová, et al., 2016).

We acknowledge that the results discussed so far are constrained to the set of species modeled (74 out of 193) which could be a biased subset of the bird community, and that there are methods to account for lack of data in some species (Devarajan, et al., 2020). But rather than assume certain parameters for the whole community, we decided to model each species separately and use only those for which there were enough observations. In fact, we encountered various species that did not behave as expected, given their dependency on forests. For example, the Scaled Antpitta (*Grallaria guatimalensis*), an understory bird associated with forests increased its occupancy at coffee plantations according to our models, whereas Azara’s Spinetail (*Synallaxis azarae*), another understory bird not associated with forests decreases its occupancy at coffee plantations. These exceptions to the general expectation enrich our vision as to how agroecosystems impact communities and allow us to anticipate the consequences of maintaining an agroecosystem dominated by sun-exposed coffee plantations in Colombia. Following our results, we predict that non-forest dependent birds, especially from the warmer lowlands, will continue to expand the upper limits of their distribution towards higher lands on the Andean mountains, further contributing to biodiversity homogenization.

We offer a potential strategy to improve forest-dependent bird occupancy in coffee agroecosystems in Jardín. Increasing FHD and CH should increase occupancy of several forest-dependent birds. This could be achieved through various strategies such as mixed cultivations (shrubs and trees), shaded-coffee plantations (coffee and trees), alternative ways to rejuvenate coffee plots in order to not have a big impact during every plant renovation cycle (light pruning) and implementing small conservation plots in sun-exposed coffee plantations (land sparing). Having coffee plantations with a shade that is composed of multiple layers at different heights will most certainly increase alpha as well as beta diversity. These strategies are not exclusive and could be mixed to minimize potential negative impacts on the farmers’ economy. Monitoring programs at coffee plantations that track dynamics in bird communities associated with these different practices is eventually the best way to evaluate their success (Doxa, et al., 2012).

## Acknowledgments

The authors would like to thank the “Our Coffee Our Birds” project in which this manuscript is part, also to Alejandro Suarez and José Castaño for their dedicated field work over several years, contributing significantly to the project’s data collection. We are very grateful to the Nespresso AAA Program for their funding, which supported these biodiversity monitoring efforts, and for their commitment to understanding the link between biodiversity and farmers’ quality of life. Also, thanks to The Cornell Lab of Ornithology for their ongoing pursuit of innovative approaches to studying, understanding, and conserving birds globally. Finally, we give our gratitude to the coffee farmers in Jardin, for their incredible interest in birds and conservation, which made this research possible.

## Notes

### Competing Interest Statement

The authors have declared no competing interest.

