## Supplementary material for "Differential Avian Responses to Coffee Farming Lead to Community Homogenization in a Working Landscape in Jardín, Colombia": Table S1

Table S1 All bird species detected in Jardin during the two sampling seasons. E, endemic; NE, near-endemic. LC, Least concern. NT, Near-threatened; VU, Vulnerable. Asterisk depicts the species that were modeled.

| **Order** | **Family** | **Scientific Name** | **English Common Name** | **Endemism** | **UICN Category** | **Forest Dependency** | **Modeled species** |
| --- | --- | --- | --- | --- | --- | --- | --- |
| Galliformes | Cracidae | *Ortalis columbiana* | Colombian Chachalaca | E | LC | none | * |
| Galliformes | Odontophoridae | *Odontophorus hyperythrus* | Chestnut Wood-Quail | E | LC | yes |  |
| Columbiformes | Columbidae | *Patagioenas fasciata* | Band-tailed Pigeon | - | LC | yes |  |
| Columbiformes | Columbidae | *Patagioenas cayennensis* | Pale-vented Pigeon | - | LC | yes |  |
| Columbiformes | Columbidae | *Patagioenas subvinacea* | Ruddy Pigeon | - | LC | yes |  |
| Columbiformes | Columbidae | *Leptotila verreauxi* | White-tipped Dove | - | LC | none | * |
| Columbiformes | Columbidae | *Zenaida auriculata* | Eared Dove | - | LC | None | * |
| Columbiformes | Columbidae | *Columbina minuta* | Plain-breasted Ground Dove | - | LC | None |  |
| Columbiformes | Columbidae | *Columbina talpacoti* | Ruddy Ground Dove | - | LC | none | * |
| Cuculiformes | Cuculidae | *Crotophaga ani* | Smooth-billed Ani | - | LC | none | * |
| Cuculiformes | Cuculidae | *Piaya cayana* | Squirrel Cuckoo | - | LC | yes | * |
| Steatornithiformes | Steatornithidae | *Steatornis caripensis* | Oilbird | - | LC | yes |  |
| Nyctibiiformes | Nyctibiidae | *Nyctibius griseus* | Common Potoo | - | LC | yes |  |
| Caprimulgiformes | Caprimulgidae | *Nyctidromus albicollis* | Common Pauraque | - | LC | yes |  |
| Apodiformes | Apodidae | *Streptoprocne rutila* | Chestnut-collared Swift | - | LC | yes |  |
| Apodiformes | Apodidae | *Streptoprocne zonaris* | White-collared Swift | - | LC | yes |  |
| Apodiformes | Apodidae | *Panyptila cayennensis* | Lesser Swallow-tailed Swift | - | LC | yes |  |
| Apodiformes | Trochilidae | *Florisuga mellivora* | White-necked Jacobin | - | LC | yes |  |
| Apodiformes | Trochilidae | *Phaethornis guy* | Green Hermit | - | LC | yes | * |
| Apodiformes | Trochilidae | *Colibri cyanotus* | Green Violetear | - | LC | none | * |
| Apodiformes | Trochilidae | *Colibri coruscans* | Sparkling Violetear | - | LC | None |  |
| Apodiformes | Trochilidae | *Anthracothorax nigricollis* | Black-throated Mango | - | LC | none | * |
| Apodiformes | Trochilidae | *Adelomyia melanogenys* | Speckled Hummingbird | - | LC | yes |  |
| Apodiformes | Trochilidae | *Aglaiocercus kingii* | Long-tailed Sylph | - | LC | None |  |
| Apodiformes | Trochilidae | *Haplophaedia aureliae* | Greenish Puffleg | NE | LC | yes |  |
| Apodiformes | Trochilidae | *Coeligena coeligena* | Bronzy Inca | - | LC | yes |  |
| Apodiformes | Trochilidae | *Ocreatus underwoodii* | Booted Racket-tail | - | LC | yes |  |
| Apodiformes | Trochilidae | *Chaetocercus mulsant* | White-bellied Woodstar | - | LC | yes |  |
| Apodiformes | Trochilidae | *Chlorostilbon melanorhynchus* | Western Emerald | NE | LC | yes | * |
| Apodiformes | Trochilidae | *Saucerottia saucerottei* | Steely-vented Hummingbird | - | LC | yes | * |
| Apodiformes | Trochilidae | *Amazilia tzacatl* | Rufous-tailed Hummingbird | - | LC | none | * |
| Apodiformes | Trochilidae | *Uranomitra franciae* | Andean Emerald | - | LC | yes | * |
| Gruiformes | Rallidae | *Pardirallus nigricans* | Blackish Rail | - | LC | None |  |
| Charadriiformes | Charadriidae | *Vanellus chilensis* | Southern Lapwing | - | LC | None |  |
| Pelecaniformes | Ardeidae | *Bubulcus ibis* | Cattle Egret | - | LC | none |  |
| Cathartiformes | Cathartidae | *Coragyps atratus* | Black Vulture | - | LC | none |  |
| Cathartiformes | Cathartidae | *Cathartes aura* | Turkey Vulture | - | LC | None |  |
| Accipitriformes | Accipitridae | *Chondrohierax uncinatus* | Hook-billed Kite | - | LC | yes |  |
| Accipitriformes | Accipitridae | *Spizaetus tyrannus* | Black Hawk-Eagle | - | LC | yes |  |
| Accipitriformes | Accipitridae | *Harpagus bidentatus* | Double-toothed Kite | - | LC | yes |  |
| Accipitriformes | Accipitridae | *Rupornis magnirostris* | Roadside Hawk | - | LC | none |  |
| Accipitriformes | Accipitridae | *Geranoaetus albicaudatus* | White-tailed Hawk | - | LC | None |  |
| Accipitriformes | Accipitridae | *Buteo brachyurus* | Short-tailed Hawk | - | LC | yes |  |
| Strigiformes | Strigidae | *Megascops choliba* | Tropical Screech-Owl | - | LC | yes |  |
| Coraciiformes | Momotidae | *Momotus aequatorialis* | Andean Motmot | - | LC | yes | * |
| Coraciiformes | Alcedinidae | *Megaceryle torquata* | Ringed Kingfisher | - | LC | yes |  |
| Galbuliformes | Bucconidae | *Malacoptila mystacalis* | Moustached Puffbird | - | LC | yes | * |
| Piciformes | Capitonidae | *Eubucco bourcierii* | Red-headed Barbet | - | LC | yes | * |
| Piciformes | Ramphastidae | *Aulacorhynchus albivitta* | Emerald Toucanet | - | LC | yes | * |
| Piciformes | Ramphastidae | *Aulacorhynchus haematopygus* | Crimson-rumped Toucanet | NE | LC | yes |  |
| Piciformes | Picidae | *Picumnus olivaceus* | Olivaceous Piculet | - | LC | yes |  |
| Piciformes | Picidae | *Picumnus granadensis* | Grayish Piculet | E | LC | yes | * |
| Piciformes | Picidae | *Melanerpes formicivorus* | Acorn Woodpecker | - | LC | yes | * |
| Piciformes | Picidae | *Melanerpes rubricapillus* | Red-crowned Woodpecker | - | LC | none | * |
| Piciformes | Picidae | *Dryobates fumigatus* | Smoky-brown Woodpecker | - | LC | yes |  |
| Piciformes | Picidae | *Dryocopus lineatus* | Lineated Woodpecker | - | LC | none | * |
| Piciformes | Picidae | *Colaptes rubiginosus* | Golden-olive Woodpecker | - | LC | yes |  |
| Piciformes | Picidae | *Colaptes punctigula* | Spot-breasted Woodpecker | - | LC | yes |  |
| Falconiformes | Falconidae | *Herpetotheres cachinnans* | Laughing Falcon | - | LC | None |  |
| Falconiformes | Falconidae | *Caracara plancus* | Crested Caracara | - | LC | None |  |
| Falconiformes | Falconidae | *Milvago chimachima* | Yellow-headed Caracara | - | LC | none | * |
| Psittaciformes | Psittacidae | *Pionus chalcopterus* | Bronze-winged Parrot | NE | LC | yes |  |
| Psittaciformes | Psittacidae | *Ognorhynchus icterotis* | Yellow-eared Parrot | NE | VU | yes |  |
| Psittaciformes | Psittacidae | *Psittacara wagleri* | Scarlet-fronted Parakeet | - | NT | yes |  |
| Passeriformes | Thamnophilidae | *Thamnophilus multistriatus* | Bar-crested Antshrike | NE | LC | yes | * |
| Passeriformes | Thamnophilidae | *Dysithamnus mentalis* | Plain Antvireo | - | LC | yes |  |
| Passeriformes | Thamnophilidae | *Cercomacroides parkeri* | Parker's Antbird | E | LC | yes | * |
| Passeriformes | Grallariidae | *Grallaria guatimalensis* | Scaled Antpitta | - | LC | yes | * |
| Passeriformes | Grallariidae | *Grallaria ruficapilla* | Chestnut-crowned Antpitta | - | LC | yes |  |
| Passeriformes | Furnariidae | *Dendrocincla fuliginosa* | Plain-brown Woodcreeper | - | LC | yes |  |
| Passeriformes | Furnariidae | *Lepidocolaptes souleyetii* | Streak-headed Woodcreeper | - | LC | yes |  |
| Passeriformes | Furnariidae | *Lepidocolaptes lacrymiger* | Montane Woodcreeper | - | LC | yes |  |
| Passeriformes | Furnariidae | *Xenops rutilans* | Streaked Xenops | - | LC | yes |  |
| Passeriformes | Furnariidae | *Lochmias nematura* | Sharp-tailed Streamcreeper | - | LC | yes |  |
| Passeriformes | Furnariidae | *Anabacerthia striaticollis* | Montane Foliage-gleaner | - | LC | yes |  |
| Passeriformes | Furnariidae | *Cranioleuca erythrops* | Red-faced Spinetail | - | LC | yes |  |
| Passeriformes | Furnariidae | *Synallaxis albescens* | Pale-breasted Spinetail | - | LC | None |  |
| Passeriformes | Furnariidae | *Synallaxis azarae* | Azara's Spinetail | - | LC | none | * |
| Passeriformes | Tyrannidae | *Phyllomyias griseiceps* | Sooty-headed Tyrannulet | - | LC | yes | * |
| Passeriformes | Tyrannidae | *Phyllomyias nigrocapillus* | Black-capped Tyrannulet | - | LC | yes |  |
| Passeriformes | Tyrannidae | *Phyllomyias cinereiceps* | Ashy-headed Tyrannulet | - | LC | yes |  |
| Passeriformes | Tyrannidae | *Phyllomyias plumbeiceps* | Plumbeous-crowned Tyrannulet | - | LC | yes |  |
| Passeriformes | Tyrannidae | *Elaenia flavogaster* | Yellow-bellied Elaenia | - | LC | none | * |
| Passeriformes | Tyrannidae | *Elaenia frantzii* | Mountain Elaenia | - | LC | yes | * |
| Passeriformes | Tyrannidae | *Camptostoma obsoletum* | Southern Beardless-Tyrannulet | - | LC | none | * |
| Passeriformes | Tyrannidae | *Serpophaga cinerea* | Torrent Tyrannulet | - | LC | yes |  |
| Passeriformes | Tyrannidae | *Zimmerius chrysops* | Golden-faced Tyrannulet | - | LC | yes | * |
| Passeriformes | Tyrannidae | *Mionectes striaticollis* | Streak-necked Flycatcher | - | LC | yes |  |
| Passeriformes | Tyrannidae | *Mionectes oleagineus* | Ochre-bellied Flycatcher | - | LC | yes | * |
| Passeriformes | Tyrannidae | *Leptopogon superciliaris* | Slaty-capped Flycatcher | - | LC | yes | * |
| Passeriformes | Tyrannidae | *Leptopogon rufipectus* | Rufous-breasted Flycatcher | NE | LC | yes |  |
| Passeriformes | Tyrannidae | *Todirostrum cinereum* | Common Tody-Flycatcher | - | LC | none | * |
| Passeriformes | Tyrannidae | *Tolmomyias sulphurescens* | Yellow-olive Flycatcher | - | LC | yes |  |
| Passeriformes | Tyrannidae | *Contopus fumigatus* | Smoke-colored Pewee | - | LC | yes |  |
| Passeriformes | Tyrannidae | *Contopus virens* | Eastern Wood-Pewee | - | LC | yes |  |
| Passeriformes | Tyrannidae | *Contopus cinereus* | Tropical Pewee | - | LC | yes | * |
| Passeriformes | Tyrannidae | *Sayornis nigricans* | Black Phoebe | - | LC | none | * |
| Passeriformes | Tyrannidae | *Machetornis rixosa* | Cattle Tyrant | - | LC | None |  |
| Passeriformes | Tyrannidae | *Myiozetetes cayanensis* | Rusty-margined Flycatcher | - | LC | none | * |
| Passeriformes | Tyrannidae | *Pitangus sulphuratus* | Great Kiskadee | - | LC | none | * |
| Passeriformes | Tyrannidae | *Myiodynastes maculatus* | Streaked Flycatcher | - | LC | none | * |
| Passeriformes | Tyrannidae | *Myiodynastes hemichrysus* | Golden-bellied Flycatcher | - | LC | yes | * |
| Passeriformes | Tyrannidae | *Tyrannus melancholicus* | Tropical Kingbird | - | LC | none | * |
| Passeriformes | Tyrannidae | *Tyrannus tyrannus* | Eastern Kingbird | - | LC | None |  |
| Passeriformes | Tyrannidae | *Myiarchus tuberculifer* | Dusky-capped Flycatcher | - | LC | yes |  |
| Passeriformes | Tyrannidae | *Myiarchus apicalis* | Apical Flycatcher | E | LC | yes |  |
| Passeriformes | Tyrannidae | *Myiarchus cephalotes* | Pale-edged Flycatcher | - | LC | yes |  |
| Passeriformes | Cotingidae | *Rupicola peruvianus* | Andean Cock-of-the-rock | - | LC | yes | * |
| Passeriformes | Pipridae | *Machaeropterus striolatus* | Striolated Manakin | - | LC | yes |  |
| Passeriformes | Tityridae | *Pachyramphus versicolor* | Barred Becard | - | LC | yes |  |
| Passeriformes | Tityridae | *Pachyramphus rufus* | Cinereous Becard | - | LC | None |  |
| Passeriformes | Tityridae | *Pachyramphus polychopterus* | White-winged Becard | - | LC | yes | * |
| Passeriformes | Vireonidae | *Cyclarhis nigrirostris* | Black-billed Peppershrike | NE | LC | yes |  |
| Passeriformes | Vireonidae | *Vireo leucophrys* | Brown-capped Vireo | - | LC | yes |  |
| Passeriformes | Vireonidae | *Vireo chivi* | Chivi Vireo | - | LC | yes |  |
| Passeriformes | Vireonidae | *Vireo olivaceus* | Red-eyed Vireo | - | LC | yes |  |
| Passeriformes | Corvidae | *Cyanocorax affinis* | Black-chested Jay | NE | LC | none |  |
| Passeriformes | Corvidae | *Cyanocorax yncas* | Green Jay | - | LC | yes | * |
| Passeriformes | Hirundinidae | *Pygochelidon cyanoleuca* | Blue-and-white Swallow | - | LC | None |  |
| Passeriformes | Hirundinidae | *Stelgidopteryx ruficollis* | Southern Rough-winged Swallow | - | LC | none |  |
| Passeriformes | Troglodytidae | *Troglodytes aedon* | House Wren | - | LC | none | * |
| Passeriformes | Troglodytidae | *Pheugopedius mystacalis* | Whiskered Wren | NE | LC | yes | * |
| Passeriformes | Troglodytidae | *Cantorchilus nigricapillus* | Bay Wren | - | LC | yes |  |
| Passeriformes | Troglodytidae | *Henicorhina leucosticta* | White-breasted Wood-Wren | - | LC | yes |  |
| Passeriformes | Troglodytidae | *Henicorhina leucophrys* | Gray-breasted Wood-Wren | - | LC | yes |  |
| Passeriformes | Cinclidae | *Cinclus leucocephalus* | White-capped Dipper | - | LC | None |  |
| Passeriformes | Turdidae | *Myadestes ralloides* | Andean Solitaire | - | LC | yes | * |
| Passeriformes | Turdidae | *Catharus aurantiirostris* | Orange-billed Nightingale-Thrush | - | LC | yes | * |
| Passeriformes | Turdidae | *Catharus ustulatus* | Swainson's Thrush | - | LC | yes |  |
| Passeriformes | Turdidae | *Turdus grayi* | Clay-colored Thrush | - | LC | None | * |
| Passeriformes | Turdidae | *Turdus ignobilis* | Black-billed Thrush | - | LC | none | * |
| Passeriformes | Mimidae | *Mimus gilvus* | Tropical Mockingbird | - | LC | None |  |
| Passeriformes | Fringillidae | *Spinus xanthogastrus* | Yellow-bellied Siskin | - | LC | None |  |
| Passeriformes | Fringillidae | *Spinus psaltria* | Lesser Goldfinch | - | LC | none | * |
| Passeriformes | Fringillidae | *Euphonia laniirostris* | Thick-billed Euphonia | - | LC | yes | * |
| Passeriformes | Fringillidae | *Chlorophonia cyanocephala* | Golden-rumped Euphonia | - | LC | yes |  |
| Passeriformes | Fringillidae | *Euphonia xanthogaster* | Orange-bellied Euphonia | - | LC | yes |  |
| Passeriformes | Fringillidae | *Chlorophonia cyanea* | Blue-naped Chlorophonia | - | LC | yes |  |
| Passeriformes | Fringillidae | *Chlorophonia pyrrhophrys* | Chestnut-breasted Chlorophonia | - | LC | yes |  |
| Passeriformes | Passerellidae | *Arremonops conirostris* | Black-striped Sparrow | - | LC | yes |  |
| Passeriformes | Passerellidae | *Arremon brunneinucha* | Chestnut-capped Brushfinch | - | LC | yes | * |
| Passeriformes | Passerellidae | *Zonotrichia capensis* | Rufous-collared Sparrow | - | LC | none |  |
| Passeriformes | Passerellidae | *Atlapetes albinucha* | White-naped Brushfinch | - | LC | yes | * |
| Passeriformes | Icteridae | *Psarocolius angustifrons* | Russet-backed Oropendola | - | LC | yes | * |
| Passeriformes | Icteridae | *Psarocolius decumanus* | Crested Oropendola | - | LC | yes | * |
| Passeriformes | Icteridae | *Cacicus uropygialis* | Scarlet-rumped Cacique | - | LC | yes |  |
| Passeriformes | Icteridae | *Icterus chrysater* | Yellow-backed Oriole | - | LC | yes | * |
| Passeriformes | Icteridae | *Molothrus oryzivorus* | Giant Cowbird | - | LC | yes |  |
| Passeriformes | Icteridae | *Molothrus bonariensis* | Shiny Cowbird | - | LC | None | * |
| Passeriformes | Icteridae | *Hypopyrrhus pyrohypogaster* | Red-bellied Grackle | E | VU | yes | * |
| Passeriformes | Parulidae | *Mniotilta varia* | Black-and-white Warbler | - | LC | yes |  |
| Passeriformes | Parulidae | *Leiothlypis peregrina* | Tennessee Warbler | - | LC | yes |  |
| Passeriformes | Parulidae | *Geothlypis philadelphia* | Mourning Warbler | - | LC | None |  |
| Passeriformes | Parulidae | *Setophaga ruticilla* | American Redstart | - | LC | yes |  |
| Passeriformes | Parulidae | *Setophaga pitiayumi* | Tropical Parula | - | LC | yes | * |
| Passeriformes | Parulidae | *Setophaga fusca* | Blackburnian Warbler | - | LC | yes | * |
| Passeriformes | Parulidae | *Setophaga petechia* | Yellow Warbler | - | LC | None |  |
| Passeriformes | Parulidae | *Myiothlypis fulvicauda* | Buff-rumped Warbler | - | LC | yes |  |
| Passeriformes | Parulidae | *Myiothlypis coronata* | Russet-crowned Warbler | - | LC | yes |  |
| Passeriformes | Parulidae | *Basileuterus culicivorus* | Golden-crowned Warbler | - | LC | yes | * |
| Passeriformes | Parulidae | *Basileuterus tristriatus* | Three-striped Warbler | - | LC | yes |  |
| Passeriformes | Parulidae | *Myioborus miniatus* | Slate-throated Redstart | - | LC | yes | * |
| Passeriformes | Cardinalidae | *Piranga flava* | Hepatic Tanager | - | LC | none | * |
| Passeriformes | Thraupidae | *Sericossypha albocristata* | White-capped Tanager | - | VU | yes |  |
| Passeriformes | Thraupidae | *Hemithraupis guira* | Guira Tanager | - | LC | yes | * |
| Passeriformes | Thraupidae | *Sicalis flaveola* | Saffron Finch | - | LC | None |  |
| Passeriformes | Thraupidae | *Diglossa sittoides* | Rusty Flowerpiercer | - | LC | None |  |
| Passeriformes | Thraupidae | *Volatinia jacarina* | Blue-black Grassquit | - | LC | none | * |
| Passeriformes | Thraupidae | *Tachyphonus rufus* | White-lined Tanager | - | LC | none | * |
| Passeriformes | Thraupidae | *Ramphocelus dimidiatus* | Crimson-backed Tanager | NE | LC | none | * |
| Passeriformes | Thraupidae | *Ramphocelus flammigerus* | Flame-rumped Tanager | - | LC | none | * |
| Passeriformes | Thraupidae | *Sporophila funerea* | Thick-billed Seed-Finch | - | LC | yes |  |
| Passeriformes | Thraupidae | *Sporophila crassirostris* | Large-billed Seed-Finch | - | LC | None |  |
| Passeriformes | Thraupidae | *Sporophila intermedia* | Gray Seedeater | - | LC | None |  |
| Passeriformes | Thraupidae | *Sporophila luctuosa* | Black-and-white Seedeater | - | LC | none | * |
| Passeriformes | Thraupidae | *Sporophila nigricollis* | Yellow-bellied Seedeater | - | LC | none | * |
| Passeriformes | Thraupidae | *Saltator maximus* | Buff-throated Saltator | - | LC | yes | * |
| Passeriformes | Thraupidae | *Saltator atripennis* | Black-winged Saltator | NE | LC | yes | * |
| Passeriformes | Thraupidae | *Saltator striatipectus* | Streaked Saltator | - | LC | none | ¨* |
| Passeriformes | Thraupidae | *Coereba flaveola* | Bananaquit | - | LC | none | * |
| Passeriformes | Thraupidae | *Tiaris olivaceus* | Yellow-faced Grassquit | - | LC | none | * |
| Passeriformes | Thraupidae | *Anisognathus somptuosus* | Blue-winged Mountain-Tanager | - | LC | yes |  |
| Passeriformes | Thraupidae | *Stilpnia heinei* | Black-capped Tanager | - | LC | yes | * |
| Passeriformes | Thraupidae | *Stilpnia vitriolina* | Scrub Tanager | NE | LC | none | * |
| Passeriformes | Thraupidae | *Stilpnia cyanicollis* | Blue-necked Tanager | - | LC | none | * |
| Passeriformes | Thraupidae | *Tangara nigroviridis* | Beryl-spangled Tanager | - | LC | yes |  |
| Passeriformes | Thraupidae | *Tangara labradorides* | Metallic-green Tanager | NE | LC | yes |  |
| Passeriformes | Thraupidae | *Tangara gyrola* | Bay-headed Tanager | - | LC | yes | * |
| Passeriformes | Thraupidae | *Tangara xanthocephala* | Saffron-crowned Tanager | - | LC | yes |  |
| Passeriformes | Thraupidae | *Tangara arthus* | Golden Tanager | - | LC | yes | * |
| Passeriformes | Thraupidae | *Thraupis episcopus* | Blue-gray Tanager | - | LC | none | * |
| Passeriformes | Thraupidae | *Thraupis palmarum* | Palm Tanager | - | LC | none | * |
