## Supplementary material for "Differential Avian Responses to Coffee Farming Lead to Community Homogenization in a Working Landscape in Jardín, Colombia": Table S2

Table S2 Similarity percentages for the 74 modeled bird species.

| **Species** | **Average** | **SD** | **Average abundance coffee** | **Average abundance not_coffee** | **Cumulative contribution** | **P** | **Forest Dependency** |
| --- | --- | --- | --- | --- | --- | --- | --- |
| *Tangara arthus* | 0,006979 | 0,002183 | 0,1009 | 0,5503 | 0,04 | 0,001 | yes |
| *Coereba flaveola* | 0,005311 | 0,003555 | 0,4633 | 0,3093 | 0,07 | 0,001 | none |
| *Elaenia frantzii* | 0,005288 | 0,004685 | 0,0766 | 0,3545 | 0,101 | 0,001 | yes |
| *Synallaxis azarae* | 0,005288 | 0,003556 | 0,3003 | 0,5259 | 0,131 | 0,001 | none |
| *Hypopyrrhus pyrohypogaster* | 0,005021 | 0,004062 | 0,361 | 0,2636 | 0,16 | 0,528 | yes |
| *Myadestes ralloides* | 0,004825 | 0,004483 | 0,0672 | 0,3233 | 0,187 | 0,001 | yes |
| *Myiodynastes maculatus* | 0,004768 | 0,003955 | 0,3076 | 0,2084 | 0,215 | 0,327 | none |
| *Ramphocelus dimidiatus* | 0,004454 | 0,003747 | 0,2886 | 0,2293 | 0,24 | 0,135 | none |
| *Zonotrichia capensis* | 0,004329 | 0,000754 | 0,937 | 0,6572 | 0,265 | 0,001 | none |
| *Myioborus miniatus* | 0,004263 | 0,003602 | 0,7252 | 0,7892 | 0,289 | 0,082 | yes |
| *Piaya cayana* | 0,00424 | 0,00299 | 0,3346 | 0,2281 | 0,314 | 0,002 | yes |
| *Troglodytes aedon* | 0,004109 | 0,002776 | 0,5406 | 0,4297 | 0,337 | 0,001 | none |
| *Crotophaga ani* | 0,003862 | 0,002689 | 0,4133 | 0,4241 | 0,359 | 0,001 | none |
| *Phyllomyias griseiceps* | 0,00372 | 0,003368 | 0,2127 | 0,1584 | 0,38 | 0,186 | yes |
| *Ramphocelus flammigerus* | 0,003715 | 0,002683 | 0,6762 | 0,5351 | 0,402 | 0,001 | none |
| *Saltator striatipectus* | 0,003628 | 0,002268 | 0,5437 | 0,4112 | 0,422 | 0,001 | none |
| *Cyanocorax yncas* | 0,00355 | 0,002421 | 0,5386 | 0,4849 | 0,443 | 0,001 | yes |
| *Pitangus sulphuratus* | 0,003423 | 0,002399 | 0,4233 | 0,4008 | 0,462 | 0,002 | none |
| *Tiaris olivaceus* | 0,003297 | 0,002628 | 0,7695 | 0,6679 | 0,481 | 0,001 | none |
| *Volatinia jacarina* | 0,003296 | 0,002356 | 0,3583 | 0,3798 | 0,5 | 0,001 | none |
| *Stilpnia heinei* | 0,003243 | 0,00203 | 0,5393 | 0,6399 | 0,519 | 0,001 | yes |
| *Myiozetetes cayanensis* | 0,003222 | 0,002307 | 0,4118 | 0,3952 | 0,537 | 0,002 | none |
| *Elaenia flavogaster* | 0,002981 | 0,002769 | 0,1461 | 0,2467 | 0,554 | 0,001 | none |
| *Uranomitra franciae* | 0,002937 | 0,002076 | 0,3567 | 0,4354 | 0,571 | 0,002 | yes |
| *Thraupis episcopus* | 0,002876 | 0,00281 | 0,8576 | 0,7521 | 0,588 | 0,001 | none |
| *Tachyphonus rufus* | 0,002653 | 0,001645 | 0,4724 | 0,3955 | 0,603 | 0,001 | none |
| *Thamnophilus multistriatus* | 0,002565 | 0,001721 | 0,3046 | 0,2 | 0,617 | 0,001 | yes |
| *Melanerpes rubricapillus* | 0,002483 | 0,001657 | 0,2674 | 0,2254 | 0,632 | 0,001 | none |
| *Momotus aequatorialis* | 0,002428 | 0,001723 | 0,607 | 0,6125 | 0,645 | 0,001 | yes |
| *Malacoptila mystacalis* | 0,002384 | 0,000213 | 0,0576 | 0,2118 | 0,659 | 0,001 | yes |
| *Anthracothorax nigricollis* | 0,002355 | 0,002143 | 0,1383 | 0,1064 | 0,673 | 0,128 | none |
| *Columbina talpacoti* | 0,002306 | 0,001886 | 0,2047 | 0,1855 | 0,686 | 0,002 | none |
| *Pheugopedius mystacalis* | 0,00226 | 0,001437 | 0,4012 | 0,5213 | 0,699 | 0,001 | yes |
| *Saucerottia saucerottei* | 0,002255 | 0,002183 | 0,8563 | 0,7785 | 0,712 | 0,001 | yes |
| *Phaethornis guy* | 0,002226 | 0,001518 | 0,3297 | 0,3073 | 0,725 | 0,001 | yes |
| *Spinus psaltria* | 0,002216 | 0,001681 | 0,3588 | 0,4121 | 0,737 | 0,001 | none |
| *Melanerpes formicivorus* | 0,002189 | 0,001621 | 0,261 | 0,3345 | 0,75 | 0,001 | yes |
| *Sporophila nigricollis* | 0,002173 | 0,001567 | 0,6986 | 0,7453 | 0,762 | 0,002 | none |
| *Saltator atripennis* | 0,002072 | 0,000972 | 0,481 | 0,6132 | 0,774 | 0,001 | yes |
| *Turdus ignobilis* | 0,002058 | 0,00146 | 0,5367 | 0,5818 | 0,786 | 0,003 | none |
| *Rupicola peruvianus* | 0,002049 | 0,001782 | 0,1649 | 0,1788 | 0,797 | 0,008 | yes |
| *Leptopogon superciliaris* | 0,00204 | 0,001418 | 0,2925 | 0,355 | 0,809 | 0,001 | yes |
| *Cercomacroides parkeri* | 0,001999 | 0,000383 | 0,2984 | 0,4276 | 0,821 | 0,001 | yes |
| *Eubucco bourcierii* | 0,001841 | 0,001354 | 0,3319 | 0,4103 | 0,831 | 0,001 | yes |
| *Sporophila luctuosa* | 0,001694 | 0,001288 | 0,2192 | 0,2932 | 0,841 | 0,001 | none |
| *Grallaria ruficapilla* | 0,001682 | 0,001412 | 0,0695 | 0,1345 | 0,851 | 0,001 | yes |
| *Aulacorhynchus albivitta* | 0,001523 | 0,001054 | 0,1405 | 0,2046 | 0,859 | 0,001 | yes |
| *Ortalis columbiana* | 0,001493 | 0,001287 | 0,8994 | 0,8258 | 0,868 | 0,001 | none |
| *Stilpnia vitriolina* | 0,001443 | 0,001036 | 0,8734 | 0,8057 | 0,876 | 0,001 | none |
| *Psarocolius angustifrons* | 0,001437 | 0,001104 | 0,7227 | 0,7077 | 0,884 | 0,001 | yes |
| *Milvago chimachima* | 0,001384 | 0,000093 | 0,6416 | 0,5519 | 0,892 | 0,001 | none |
| *Grallaria guatimalensis* | 0,001347 | 0,001041 | 0,1757 | 0,1752 | 0,9 | 0,01 | yes |
| *Leptotila verreauxi* | 0,001298 | 0,000626 | 0,5091 | 0,4266 | 0,907 | 0,001 | none |
| *Myiodynastes chrysocephalus* | 0,001287 | 0,000857 | 0,3646 | 0,3695 | 0,915 | 0,001 | yes |
| *Amazilia tzacatl* | 0,001258 | 0,001063 | 0,7475 | 0,7255 | 0,922 | 0,062 | none |
| *Tyrannus melancholicus* | 0,001112 | 0,000918 | 0,8559 | 0,828 | 0,928 | 0,001 | none |
| *Todirostrum cinereum* | 0,001061 | 0,000868 | 0,1941 | 0,2127 | 0,934 | 0,001 | none |
| *Picumnus granadensis* | 0,001011 | 0,000727 | 0,2994 | 0,2776 | 0,94 | 0,001 | yes |
| *Pachyramphus polychopterus* | 0,001003 | 0,000688 | 0,2074 | 0,19 | 0,946 | 0,001 | yes |
| *Tangara gyrola* | 0,000991 | 0,000658 | 0,5375 | 0,5752 | 0,952 | 0,001 | yes |
| *Zimmerius chrysops* | 0,000953 | 0,000647 | 0,6241 | 0,5937 | 0,957 | 0,001 | yes |
| *Stilpnia cyanicollis* | 0,000881 | 0,000618 | 0,4993 | 0,4804 | 0,962 | 0,001 | none |
| *Sayornis nigricans* | 0,000875 | 0,000982 | 0,0392 | 0,0856 | 0,967 | 0,001 | none |
| *Thraupis palmarum* | 0,000847 | 0,000333 | 0,8064 | 0,7518 | 0,972 | 0,001 | none |
| *Arremon brunneinucha* | 0,000846 | 0,000573 | 0,7625 | 0,7612 | 0,977 | 0,001 | yes |
| *Icterus chrysater* | 0,000763 | 0,000675 | 0,9611 | 0,9259 | 0,981 | 0,001 | yes |
| *Atlapetes albinucha* | 0,000652 | 0,000627 | 0,8568 | 0,8464 | 0,985 | 0,001 | yes |
| *Piranga flava* | 0,000575 | 0,00043 | 0,2593 | 0,2473 | 0,988 | 0,001 | none |
| *Hemithraupis guira* | 0,000522 | 0,000572 | 0,0665 | 0,0646 | 0,991 | 0,01 | yes |
| *Colibri cyanotus* | 0,000422 | 0,000318 | 0,187 | 0,1808 | 0,994 | 0,002 | none |
| *Chlorostilbon melanorhynchus* | 0,00036 | 0,000152 | 0,7758 | 0,7526 | 0,996 | 0,001 | yes |
| *Dryocopus lineatus* | 0,000344 | 0,000293 | 0,7641 | 0,7652 | 0,998 | 0,247 | none |
| *Catharus aurantiirostris* | 0,000298 | 0,000229 | 0,3387 | 0,3424 | 0,999 | 0,002 | yes |
| *Camptostoma obsoletum* | 0,000127 | 0,000097 | 0,0761 | 0,0814 | 1 | 0,001 | none |
