## Supplementary material for "Differential Avian Responses to Coffee Farming Lead to Community Homogenization in a Working Landscape in Jardín, Colombia": Table S3

Table S3 Similarity percentages for all species detected.

| **Species** | **Average** | **SD** | **Average abundance coffee** | **Average abundance not_coffee** | **Cumulative contribution** | **P** | **Forest Dependency** |
| --- | --- | --- | --- | --- | --- | --- | --- |
| *Icterus chrysater* | 0,0255 | 0,019029 | 4,207 | 2,6905 | 0,037 | 0,001 | yes |
| *Zonotrichia capensis* | 0,022333 | 0,016897 | 2,685 | 1,2381 | 0,07 | 0,003 | none |
| *Myioborus miniatus* | 0,019869 | 0,017071 | 1,333 | 2,0952 | 0,099 | 0,007 | yes |
| *Tyrannus melancholicus* | 0,018339 | 0,016953 | 1,414 | 2,0238 | 0,125 | 0,001 | none |
| *Pionus chalcopterus* | 0,01673 | 0,014406 | 1,676 | 1,1429 | 0,15 | 0,29 | yes |
| *Ramphocelus flammigerus* | 0,015696 | 0,015154 | 1,342 | 1,0476 | 0,173 | 0,352 | none |
| *Ortalis columbiana* | 0,015686 | 0,013798 | 1,748 | 1,381 | 0,196 | 0,503 | none |
| *Zimmerius chrysops* | 0,015319 | 0,015511 | 1,18 | 1,2143 | 0,218 | 0,416 | yes |
| *Psarocolius angustifrons* | 0,015095 | 0,014348 | 1,243 | 1,1905 | 0,24 | 0,413 | yes |
| *Thraupis episcopus* | 0,014967 | 0,012061 | 1,477 | 1,1667 | 0,262 | 0,123 | none |
| *Tiaris olivaceus* | 0,014416 | 0,01361 | 1,396 | 0,9048 | 0,283 | 0,488 | none |
| *Bubulcus ibis* | 0,013753 | 0,013667 | 1,225 | 0,9286 | 0,303 | 0,183 | none |
| *Rupornis magnirostris* | 0,013428 | 0,012086 | 1,523 | 1,1429 | 0,322 | 0,889 | none |
| *Cyanocorax affinis* | 0,012658 | 0,012066 | 1,27 | 0,9286 | 0,341 | 0,562 | none |
| *Atlapetes albinucha* | 0,01241 | 0,012487 | 1,144 | 0,7381 | 0,359 | 0,192 | yes |
| *Stilpnia vitriolina* | 0,0124 | 0,011075 | 1,144 | 1,1905 | 0,377 | 0,325 | none |
| *Troglodytes aedon* | 0,012119 | 0,014735 | 0,973 | 0,6429 | 0,395 | 0,72 | none |
| *Saucerottia saucerottei* | 0,011044 | 0,010283 | 1,153 | 0,6667 | 0,411 | 0,854 | yes |
| *Stilpnia heinei* | 0,010976 | 0,012458 | 0,595 | 0,9762 | 0,427 | 0,017 | yes |
| *Pheugopedius mystacalis* | 0,010321 | 0,011711 | 0,495 | 0,881 | 0,442 | 0,036 | yes |
| *Thraupis palmarum* | 0,010082 | 0,009774 | 0,955 | 0,6429 | 0,456 | 0,742 | none |
| *Momotus aequatorialis* | 0,009805 | 0,012266 | 0,432 | 0,8333 | 0,471 | 0,019 | yes |
| *Tangara gyrola* | 0,009794 | 0,010486 | 0,459 | 0,9048 | 0,485 | 0,002 | yes |
| *Coragyps atratus* | 0,009176 | 0,010198 | 0,757 | 0,4286 | 0,498 | 0,44 | none |
| *Patagioenas cayennensis* | 0,008942 | 0,010546 | 0,622 | 0,5238 | 0,511 | 0,257 | yes |
| *Turdus ignobilis* | 0,00879 | 0,009748 | 0,55 | 0,7143 | 0,524 | 0,187 | none |
| *Stelgidopteryx ruficollis* | 0,008122 | 0,010215 | 0,64 | 0,4286 | 0,536 | 0,801 | none |
| *Phaethornis guy* | 0,007878 | 0,013663 | 0,297 | 0,5714 | 0,548 | 0,035 | yes |
| *Chlorostilbon melanorhynchus* | 0,007678 | 0,0088 | 0,685 | 0,4286 | 0,559 | 0,93 | yes |
| *Tangara arthus* | 0,007536 | 0,010412 | 0,09 | 0,7381 | 0,57 | 0,001 | yes |
| *Amazilia tzacatl* | 0,007521 | 0,00895 | 0,586 | 0,4048 | 0,581 | 0,414 | none |
| *Cercomacroides parkeri* | 0,00747 | 0,010789 | 0,27 | 0,5714 | 0,592 | 0,035 | yes |
| *Saltator atripennis* | 0,007279 | 0,009046 | 0,423 | 0,5476 | 0,602 | 0,208 | yes |
| *Coereba flaveola* | 0,007209 | 0,009848 | 0,541 | 0,4048 | 0,613 | 0,775 | none |
| *Arremon brunneinucha* | 0,006843 | 0,008319 | 0,45 | 0,4286 | 0,623 | 0,248 | yes |
| *Cyanocorax yncas* | 0,006823 | 0,008739 | 0,432 | 0,4286 | 0,633 | 0,58 | yes |
| *Myiozetetes cayanensis* | 0,006681 | 0,00912 | 0,396 | 0,4524 | 0,642 | 0,319 | none |
| *Catharus aurantiirostris* | 0,006582 | 0,009702 | 0,432 | 0,381 | 0,652 | 0,631 | yes |
| *Stilpnia cyanicollis* | 0,006576 | 0,008451 | 0,216 | 0,5952 | 0,662 | 0,001 | none |
| *Dryocopus lineatus* | 0,006502 | 0,007483 | 0,523 | 0,3571 | 0,671 | 0,924 | none |
| *Leptotila verreauxi* | 0,006501 | 0,008976 | 0,468 | 0,3095 | 0,68 | 0,733 | none |
| *Saltator striatipectus* | 0,006325 | 0,008855 | 0,45 | 0,3571 | 0,69 | 0,52 | none |
| *Sporophila nigricollis* | 0,006299 | 0,007794 | 0,45 | 0,3571 | 0,699 | 0,541 | none |
| *Eubucco bourcierii* | 0,006289 | 0,00894 | 0,252 | 0,5 | 0,708 | 0,019 | yes |
| *Volatinia jacarina* | 0,006161 | 0,009989 | 0,333 | 0,4048 | 0,717 | 0,261 | none |
| *Hypopyrrhus pyrohypogaster* | 0,005537 | 0,008506 | 0,342 | 0,2857 | 0,725 | 0,603 | yes |
| *Myiodynastes chrysocephalus* | 0,005173 | 0,007966 | 0,234 | 0,3571 | 0,733 | 0,133 | yes |
| *Colaptes rubiginosus* | 0,004957 | 0,006146 | 0,396 | 0,2857 | 0,74 | 0,833 | yes |
| *Pitangus sulphuratus* | 0,004707 | 0,007662 | 0,342 | 0,2381 | 0,747 | 0,81 | none |
| *Synallaxis azarae* | 0,004691 | 0,007913 | 0,18 | 0,3571 | 0,754 | 0,065 | none |
| *Milvago chimachima* | 0,004627 | 0,006748 | 0,297 | 0,2619 | 0,76 | 0,424 | none |
| *Psittacara wagleri* | 0,004609 | 0,007276 | 0,18 | 0,381 | 0,767 | 0,062 | yes |
| *Grallaria ruficapilla* | 0,004515 | 0,014159 | 0,054 | 0,3571 | 0,774 | 0,006 | yes |
| *Spinus psaltria* | 0,004467 | 0,007694 | 0,243 | 0,2857 | 0,78 | 0,306 | none |
| *Tachyphonus rufus* | 0,004344 | 0,007154 | 0,261 | 0,2857 | 0,786 | 0,396 | none |
| *Crotophaga ani* | 0,004305 | 0,00702 | 0,315 | 0,1905 | 0,793 | 0,91 | none |
| *Sporophila luctuosa* | 0,004265 | 0,007973 | 0,207 | 0,2857 | 0,799 | 0,282 | none |
| *Uranomitra franciae* | 0,004241 | 0,007407 | 0,216 | 0,2619 | 0,805 | 0,244 | yes |
| *Patagioenas fasciata* | 0,004219 | 0,007519 | 0,072 | 0,381 | 0,811 | 0,001 | yes |
| *Myadestes ralloides* | 0,004099 | 0,00916 | 0,018 | 0,381 | 0,817 | 0,001 | yes |
| *Piaya cayana* | 0,00399 | 0,006973 | 0,306 | 0,1667 | 0,823 | 0,915 | yes |
| *Leptopogon superciliaris* | 0,003884 | 0,007477 | 0,108 | 0,3333 | 0,829 | 0,005 | yes |
| *Thamnophilus multistriatus* | 0,003881 | 0,007335 | 0,288 | 0,1429 | 0,835 | 0,912 | yes |
| *Vanellus chilensis* | 0,003797 | 0,007109 | 0,063 | 0,3333 | 0,84 | 0,004 | None |
| *Vireo leucophrys* | 0,003776 | 0,00961 | 0,018 | 0,3333 | 0,846 | 0,001 | yes |
| *Elaenia frantzii* | 0,003738 | 0,006826 | 0,054 | 0,3571 | 0,851 | 0,001 | yes |
| *Melanerpes formicivorus* | 0,003708 | 0,007554 | 0,207 | 0,2381 | 0,856 | 0,424 | yes |
| *Streptoprocne zonaris* | 0,003618 | 0,006178 | 0,243 | 0,1667 | 0,862 | 0,861 | yes |
| *Grallaria guatimalensis* | 0,003283 | 0,007972 | 0,198 | 0,1429 | 0,866 | 0,671 | yes |
| *Pachyramphus polychopterus* | 0,00312 | 0,006633 | 0,171 | 0,1905 | 0,871 | 0,373 | yes |
| *Pygochelidon cyanoleuca* | 0,002978 | 0,006226 | 0,243 | 0,0714 | 0,875 | 0,977 | None |
| *Todirostrum cinereum* | 0,00266 | 0,005507 | 0,153 | 0,1667 | 0,879 | 0,509 | none |
| *Streptoprocne rutila* | 0,002538 | 0,006124 | 0,198 | 0,0476 | 0,883 | 0,962 | yes |
| *Sayornis nigricans* | 0,002532 | 0,008886 | 0,054 | 0,1905 | 0,887 | 0,049 | none |
| *Cathartes aura* | 0,002388 | 0,005501 | 0,18 | 0,0714 | 0,89 | 0,916 | None |
| *Elaenia flavogaster* | 0,002297 | 0,004967 | 0,045 | 0,2381 | 0,893 | 0,009 | none |
| *Ramphocelus dimidiatus* | 0,002261 | 0,004885 | 0,108 | 0,1667 | 0,897 | 0,236 | none |
| *Aulacorhynchus albivitta* | 0,002254 | 0,005751 | 0,099 | 0,119 | 0,9 | 0,427 | yes |
| *Rupicola peruvianus* | 0,002191 | 0,006184 | 0,036 | 0,1667 | 0,903 | 0,022 | yes |
| *Columbina talpacoti* | 0,002121 | 0,005008 | 0,099 | 0,1667 | 0,906 | 0,24 | none |
| *Myiodynastes maculatus* | 0,002114 | 0,004791 | 0,108 | 0,119 | 0,909 | 0,325 | none |
| *Melanerpes rubricapillus* | 0,001981 | 0,004609 | 0,063 | 0,1905 | 0,912 | 0,08 | none |
| *Malacoptila mystacalis* | 0,001959 | 0,00557 | 0,027 | 0,1429 | 0,915 | 0,026 | yes |
| *Herpetotheres cachinnans* | 0,001939 | 0,004812 | 0,126 | 0,0714 | 0,918 | 0,793 | None |
| *Picumnus granadensis* | 0,001878 | 0,004305 | 0,081 | 0,1429 | 0,921 | 0,176 | yes |
| *Ognorhynchus icterotis* | 0,001851 | 0,005039 | 0,063 | 0,119 | 0,923 | 0,159 | yes |
| *Synallaxis albescens* | 0,001526 | 0,004055 | 0,072 | 0,0952 | 0,926 | 0,277 | None |
| *Aulacorhynchus haematopygus* | 0,00148 | 0,003684 | 0,108 | 0,0476 | 0,928 | 0,869 | yes |
| *Setophaga fusca* | 0,001417 | 0,004971 | 0,054 | 0,0714 | 0,93 | 0,235 | yes |
| *Anthracothorax nigricollis* | 0,001361 | 0,004468 | 0,126 | 0,0238 | 0,932 | 0,95 | none |
| *Ocreatus underwoodii* | 0,001327 | 0,005619 | 0 | 0,119 | 0,934 | 0,019 | yes |
| *Phyllomyias griseiceps* | 0,001305 | 0,003513 | 0,09 | 0,0714 | 0,936 | 0,607 | yes |
| *Piranga flava* | 0,001288 | 0,00382 | 0,117 | 0,0238 | 0,938 | 0,973 | none |
| *Setophaga pitiayumi* | 0,001275 | 0,003511 | 0,063 | 0,0714 | 0,939 | 0,287 | yes |
| *Psarocolius decumanus* | 0,001263 | 0,003724 | 0,063 | 0,0714 | 0,941 | 0,409 | yes |
| *Mionectes oleagineus* | 0,00126 | 0,003754 | 0,018 | 0,119 | 0,943 | 0,014 | yes |
| *Colibri cyanotus* | 0,001095 | 0,003835 | 0,108 | 0 | 0,945 | 0,977 | none |
| *Myiarchus cephalotes* | 0,001088 | 0,003034 | 0 | 0,119 | 0,946 | 0,004 | yes |
| *Molothrus bonariensis* | 0,001076 | 0,003014 | 0,054 | 0,0714 | 0,948 | 0,442 | None |
| *Hemithraupis guira* | 0,001072 | 0,004267 | 0,063 | 0,0714 | 0,949 | 0,444 | yes |
| *Contopus cinereus* | 0,00106 | 0,003227 | 0,036 | 0,0714 | 0,951 | 0,1 | yes |
| *Zenaida auriculata* | 0,001042 | 0,003477 | 0,063 | 0,0476 | 0,952 | 0,72 | None |
| *Molothrus oryzivorus* | 0,001008 | 0,003102 | 0,081 | 0,0238 | 0,954 | 0,914 | yes |
| *Contopus fumigatus* | 0,001007 | 0,004901 | 0 | 0,0952 | 0,955 | 0,048 | yes |
| *Camptostoma obsoletum* | 0,000986 | 0,003608 | 0,045 | 0,0714 | 0,957 | 0,342 | none |
| *Tangara labradorides* | 0,00098 | 0,00377 | 0 | 0,0952 | 0,958 | 0,013 | yes |
| *Basileuterus culicivorus* | 0,000957 | 0,00334 | 0,036 | 0,0476 | 0,96 | 0,227 | yes |
| *Cinclus leucocephalus* | 0,000946 | 0,004487 | 0 | 0,0714 | 0,961 | 0,035 | None |
| *Myiothlypis fulvicauda* | 0,000923 | 0,004212 | 0,045 | 0,0476 | 0,962 | 0,38 | yes |
| *Sporophila funerea* | 0,000788 | 0,00388 | 0 | 0,0952 | 0,964 | 0,061 | yes |
| *Anisognathus somptuosus* | 0,000781 | 0,003786 | 0 | 0,0714 | 0,965 | 0,062 | yes |
| *Euphonia laniirostris* | 0,000772 | 0,002628 | 0,063 | 0,0238 | 0,966 | 0,881 | yes |
| *Saltator maximus* | 0,000736 | 0,002578 | 0,036 | 0,0476 | 0,967 | 0,454 | yes |
| *Phyllomyias nigrocapillus* | 0,000717 | 0,002814 | 0,018 | 0,0476 | 0,968 | 0,131 | yes |
| *Chlorophonia cyanocephala* | 0,000686 | 0,003161 | 0,054 | 0,0238 | 0,969 | 0,747 | yes |
| *Chlorophonia cyanea* | 0,000651 | 0,003045 | 0 | 0,0714 | 0,97 | 0,058 | yes |
| *Caracara plancus* | 0,000627 | 0,002309 | 0,027 | 0,0476 | 0,971 | 0,267 | None |
| *Turdus grayi* | 0,000605 | 0,002558 | 0,045 | 0,0238 | 0,972 | 0,724 | None |
| *Myiothlypis coronata* | 0,000557 | 0,002551 | 0 | 0,0476 | 0,972 | 0,051 | yes |
| *Chondrohierax uncinatus* | 0,000518 | 0,002822 | 0,045 | 0 | 0,973 | 0,779 | yes |
| *Mimus gilvus* | 0,000514 | 0,002163 | 0,036 | 0,0238 | 0,974 | 0,664 | None |
| *Vireo chivi* | 0,000494 | 0,002078 | 0,009 | 0,0476 | 0,975 | 0,118 | yes |
| *Henicorhina leucophrys* | 0,000493 | 0,002257 | 0 | 0,0476 | 0,975 | 0,046 | yes |
| *Serpophaga cinerea* | 0,000486 | 0,002298 | 0 | 0,0476 | 0,976 | 0,058 | yes |
| *Tolmomyias sulphurescens* | 0,000481 | 0,002471 | 0,018 | 0,0238 | 0,977 | 0,283 | yes |
| *Nyctibius griseus* | 0,00047 | 0,003064 | 0,054 | 0 | 0,978 | 0,758 | yes |
| *Coeligena coeligena* | 0,000465 | 0,003007 | 0 | 0,0476 | 0,978 | 0,184 | yes |
| *Tangara nigroviridis* | 0,000465 | 0,003007 | 0 | 0,0476 | 0,979 | 0,184 | yes |
| *Xenops rutilans* | 0,000465 | 0,003007 | 0 | 0,0476 | 0,98 | 0,184 | yes |
| *Patagioenas subvinacea* | 0,000458 | 0,002072 | 0 | 0,0476 | 0,98 | 0,057 | yes |
| *Dryobates fumigatus* | 0,000443 | 0,001863 | 0,009 | 0,0476 | 0,981 | 0,187 | yes |
| *Contopus virens* | 0,00042 | 0,002344 | 0,009 | 0,0238 | 0,981 | 0,292 | yes |
| *Tangara xanthocephala* | 0,000419 | 0,001891 | 0 | 0,0476 | 0,982 | 0,057 | yes |
| *Adelomyia melanogenys* | 0,000409 | 0,002029 | 0,018 | 0,0238 | 0,983 | 0,508 | yes |
| *Megaceryle torquata* | 0,000398 | 0,002065 | 0,018 | 0,0238 | 0,983 | 0,501 | yes |
| *Phyllomyias plumbeiceps* | 0,000387 | 0,002157 | 0,009 | 0,0238 | 0,984 | 0,215 | yes |
| *Lochmias nematura* | 0,000385 | 0,002145 | 0,009 | 0,0238 | 0,984 | 0,223 | yes |
| *Lepidocolaptes lacrymiger* | 0,000385 | 0,001913 | 0,018 | 0,0238 | 0,985 | 0,371 | yes |
| *Geothlypis philadelphia* | 0,00038 | 0,002125 | 0,009 | 0,0238 | 0,986 | 0,209 | None |
| *Arremonops conirostris* | 0,000375 | 0,00242 | 0 | 0,0476 | 0,986 | 0,213 | yes |
| *Phyllomyias cinereiceps* | 0,000373 | 0,002083 | 0,009 | 0,0238 | 0,987 | 0,223 | yes |
| *Cacicus uropygialis* | 0,000368 | 0,001935 | 0,036 | 0 | 0,987 | 0,818 | yes |
| *Colibri coruscans* | 0,000335 | 0,001856 | 0,009 | 0,0238 | 0,988 | 0,214 | None |
| *Chlorophonia pyrrhophrys* | 0,000333 | 0,002178 | 0 | 0,0238 | 0,988 | 0,179 | yes |
| *Leptopogon rufipectus* | 0,00031 | 0,002024 | 0 | 0,0238 | 0,989 | 0,165 | yes |
| *Mionectes striaticollis* | 0,00031 | 0,002024 | 0 | 0,0238 | 0,989 | 0,165 | yes |
| *Geranoaetus albicaudatus* | 0,000302 | 0,001969 | 0 | 0,0238 | 0,989 | 0,194 | None |
| *Chaetocercus mulsant* | 0,000297 | 0,001557 | 0,036 | 0 | 0,99 | 0,856 | yes |
| *Sporophila intermedia* | 0,000289 | 0,00163 | 0,009 | 0,0238 | 0,99 | 0,306 | None |
| *Pachyramphus versicolor* | 0,000287 | 0,001867 | 0 | 0,0238 | 0,991 | 0,163 | yes |
| *Panyptila cayennensis* | 0,000267 | 0,001734 | 0 | 0,0238 | 0,991 | 0,203 | yes |
| *Dysithamnus mentalis* | 0,000264 | 0,001714 | 0 | 0,0238 | 0,991 | 0,21 | yes |
| *Columbina minuta* | 0,000252 | 0,001637 | 0 | 0,0238 | 0,992 | 0,196 | None |
| *Basileuterus tristriatus* | 0,000247 | 0,001602 | 0 | 0,0238 | 0,992 | 0,203 | yes |
| *Cyclarhis nigrirostris* | 0,000247 | 0,001602 | 0 | 0,0238 | 0,993 | 0,203 | yes |
| *Henicorhina leucosticta* | 0,000239 | 0,001551 | 0 | 0,0238 | 0,993 | 0,2 | yes |
| *Pardirallus nigricans* | 0,000235 | 0,001519 | 0 | 0,0238 | 0,993 | 0,184 | None |
| *Steatornis caripensis* | 0,000235 | 0,001519 | 0 | 0,0238 | 0,994 | 0,201 | yes |
| *Aglaiocercus kingii* | 0,000232 | 0,001504 | 0 | 0,0238 | 0,994 | 0,184 | None |
| *Tyrannus tyrannus* | 0,000211 | 0,001365 | 0 | 0,0238 | 0,994 | 0,191 | None |
| *Anabacerthia striaticollis* | 0,000206 | 0,001329 | 0 | 0,0238 | 0,995 | 0,201 | yes |
| *Euphonia xanthogaster* | 0,000206 | 0,001329 | 0 | 0,0238 | 0,995 | 0,201 | yes |
| *Haplophaedia aureliae* | 0,000206 | 0,001329 | 0 | 0,0238 | 0,995 | 0,201 | yes |
| *Nyctidromus albicollis* | 0,000187 | 0,001997 | 0,018 | 0 | 0,995 | 0,499 | yes |
| *Mniotilta varia* | 0,000183 | 0,001368 | 0,018 | 0 | 0,996 | 0,652 | yes |
| *Myiarchus apicalis* | 0,000178 | 0,001332 | 0,018 | 0 | 0,996 | 0,669 | yes |
| *Picumnus olivaceus* | 0,000176 | 0,001136 | 0 | 0,0238 | 0,996 | 0,229 | yes |
| *Setophaga ruticilla* | 0,000165 | 0,001239 | 0,018 | 0 | 0,996 | 0,669 | yes |
| *Colaptes punctigula* | 0,000156 | 0,001007 | 0 | 0,0238 | 0,997 | 0,217 | yes |
| *Vireo olivaceus* | 0,000156 | 0,001007 | 0 | 0,0238 | 0,997 | 0,217 | yes |
| *Lepidocolaptes souleyetii* | 0,000154 | 0,001185 | 0,018 | 0 | 0,997 | 0,646 | yes |
| *Sporophila crassirostris* | 0,000149 | 0,001581 | 0,018 | 0 | 0,997 | 0,553 | None |
| *Catharus ustulatus* | 0,000148 | 0,001568 | 0,018 | 0 | 0,998 | 0,526 | yes |
| *Machetornis rixosa* | 0,000112 | 0,001198 | 0,009 | 0 | 0,998 | 0,491 | None |
| *Sicalis flaveola* | 0,000109 | 0,001167 | 0,009 | 0 | 0,998 | 0,483 | None |
| *Harpagus bidentatus* | 0,000108 | 0,001152 | 0,009 | 0 | 0,998 | 0,496 | yes |
| *Cantorchilus nigricapillus* | 0,000101 | 0,001082 | 0,009 | 0 | 0,998 | 0,501 | yes |
| *Setophaga petechia* | 0,000101 | 0,001082 | 0,009 | 0 | 0,998 | 0,495 | None |
| *Odontophorus hyperythrus* | 0,000096 | 0,001021 | 0,009 | 0 | 0,998 | 0,494 | yes |
| *Pachyramphus rufus* | 0,000096 | 0,001021 | 0,009 | 0 | 0,999 | 0,491 | None |
| *Spinus xanthogastrus* | 0,000095 | 0,001009 | 0,009 | 0 | 0,999 | 0,501 | None |
| *Buteo brachyurus* | 0,000092 | 0,000977 | 0,009 | 0 | 0,999 | 0,498 | yes |
| *Megascops choliba* | 0,000091 | 0,000966 | 0,009 | 0 | 0,999 | 0,517 | yes |
| *Sericossypha albocristata* | 0,000091 | 0,000966 | 0,009 | 0 | 0,999 | 0,5 | yes |
| *Spizaetus tyrannus* | 0,000091 | 0,000966 | 0,009 | 0 | 0,999 | 0,5 | yes |
| *Myiarchus tuberculifer* | 0,000086 | 0,000918 | 0,009 | 0 | 0,999 | 0,527 | yes |
| *Machaeropterus striolatus* | 0,000084 | 0,000891 | 0,009 | 0 | 1 | 0,506 | yes |
| *Diglossa sittoides* | 0,000079 | 0,000834 | 0,009 | 0 | 1 | 0,528 | None |
| *Cranioleuca erythrops* | 0,000078 | 0,000826 | 0,009 | 0 | 1 | 0,534 | yes |
| *Leiothlypis peregrina* | 0,000074 | 0,000784 | 0,009 | 0 | 1 | 0,522 | yes |
| *Dendrocincla fuliginosa* | 0,00006 | 0,000629 | 0,009 | 0 | 1 | 0,56 | yes |
| *Florisuga mellivora* | 0,00006 | 0,000629 | 0,009 | 0 | 1 | 0,56 | yes |
