## Supplementary material for "Differential Avian Responses to Coffee Farming Lead to Community Homogenization in a Working Landscape in Jardín, Colombia": Figure S1

Figure S1 Maps displaying the four covariates used in the analyses. 1) Foliage Height Diversity (FHD), 2) Canopy Height (CH), 3) Human Footprint (LHFI) and 4) Elevation (EL).

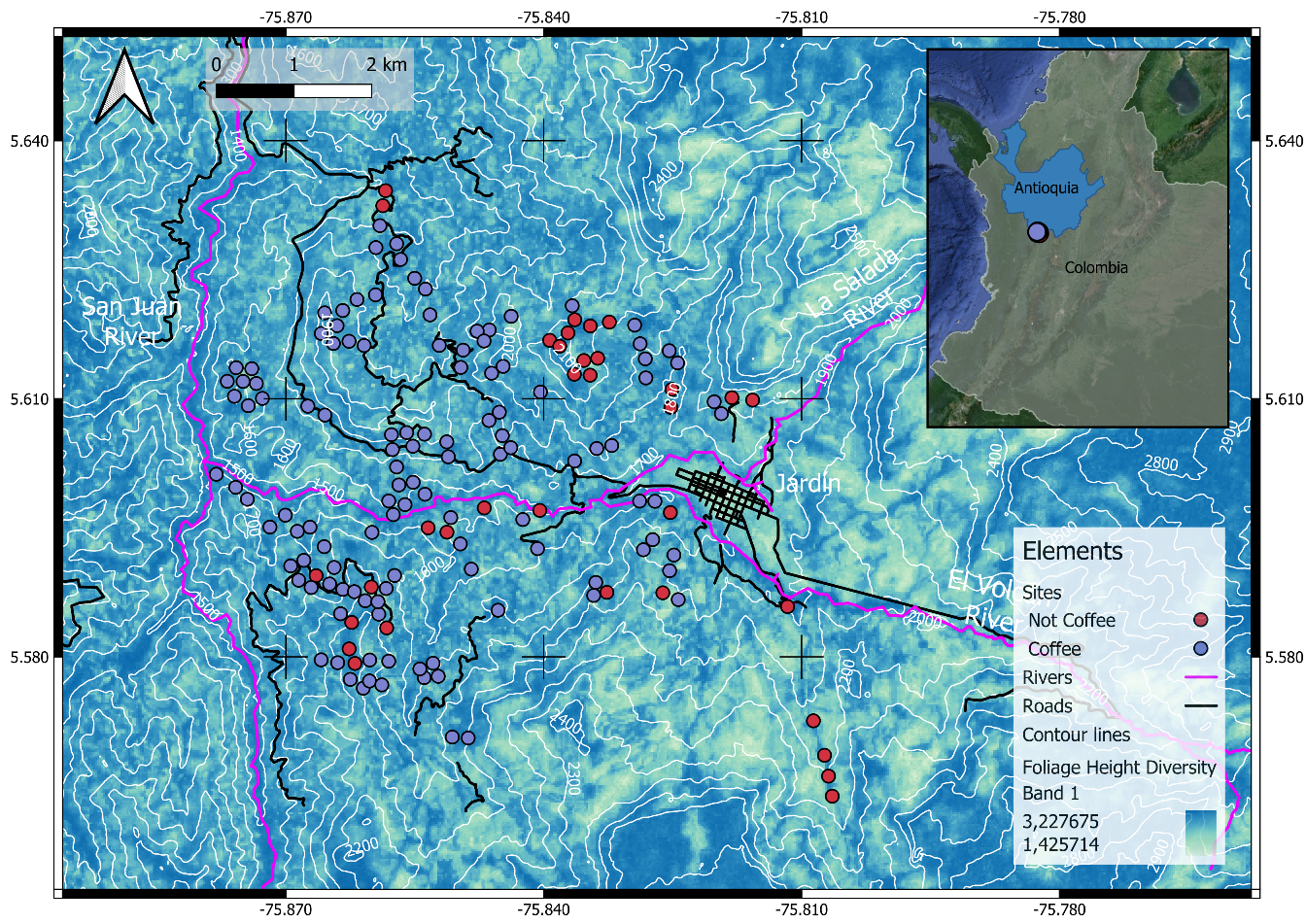

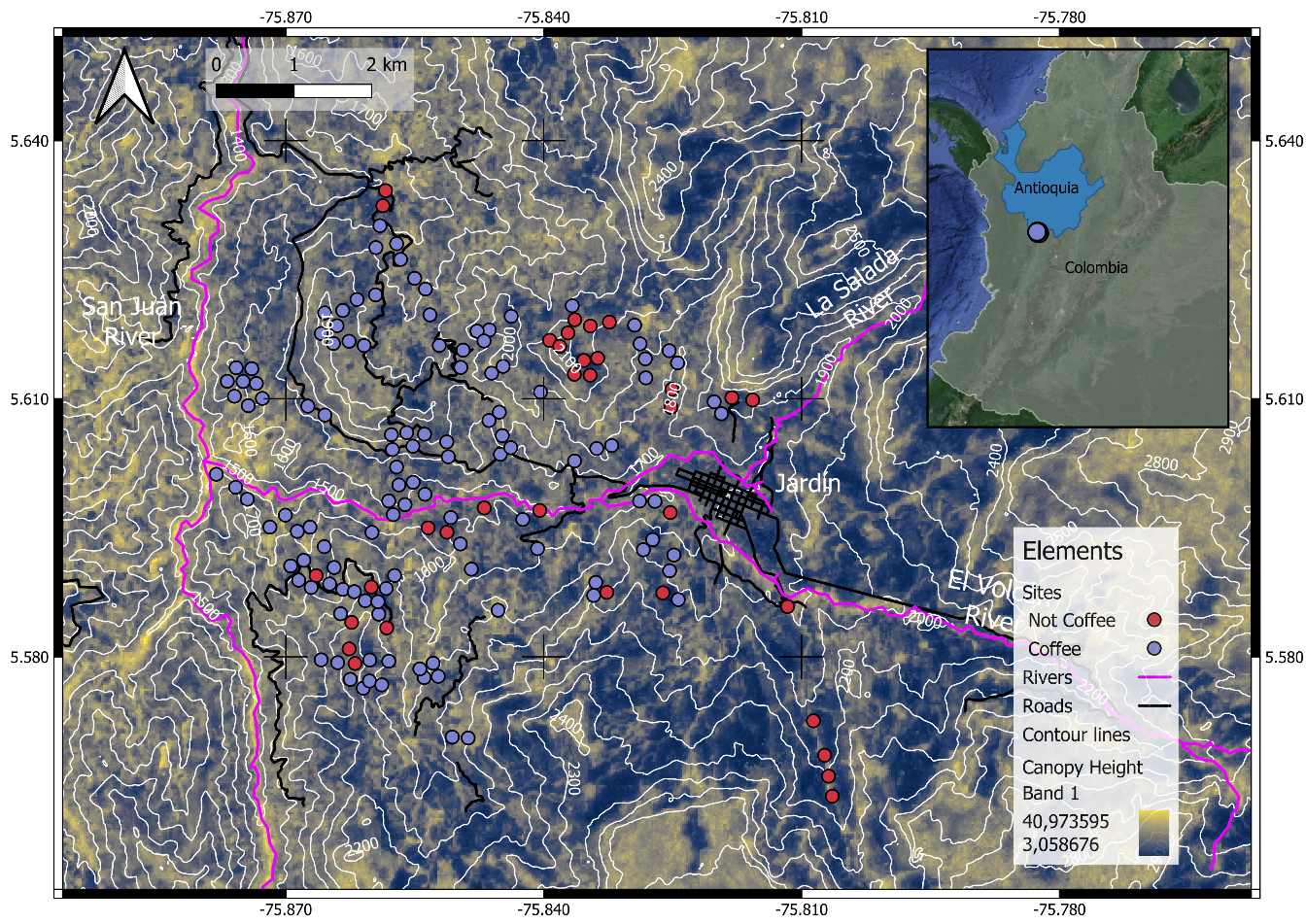

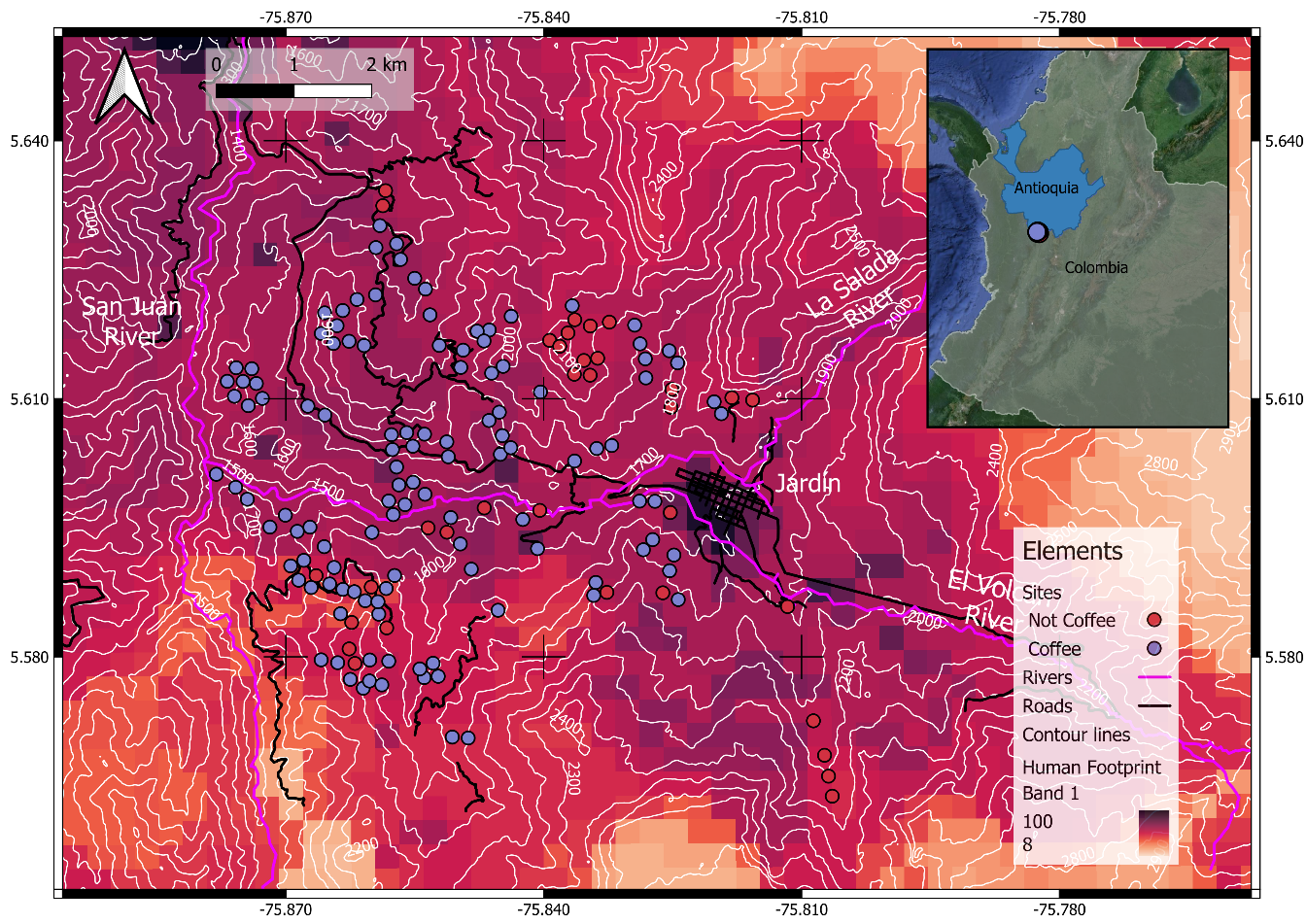

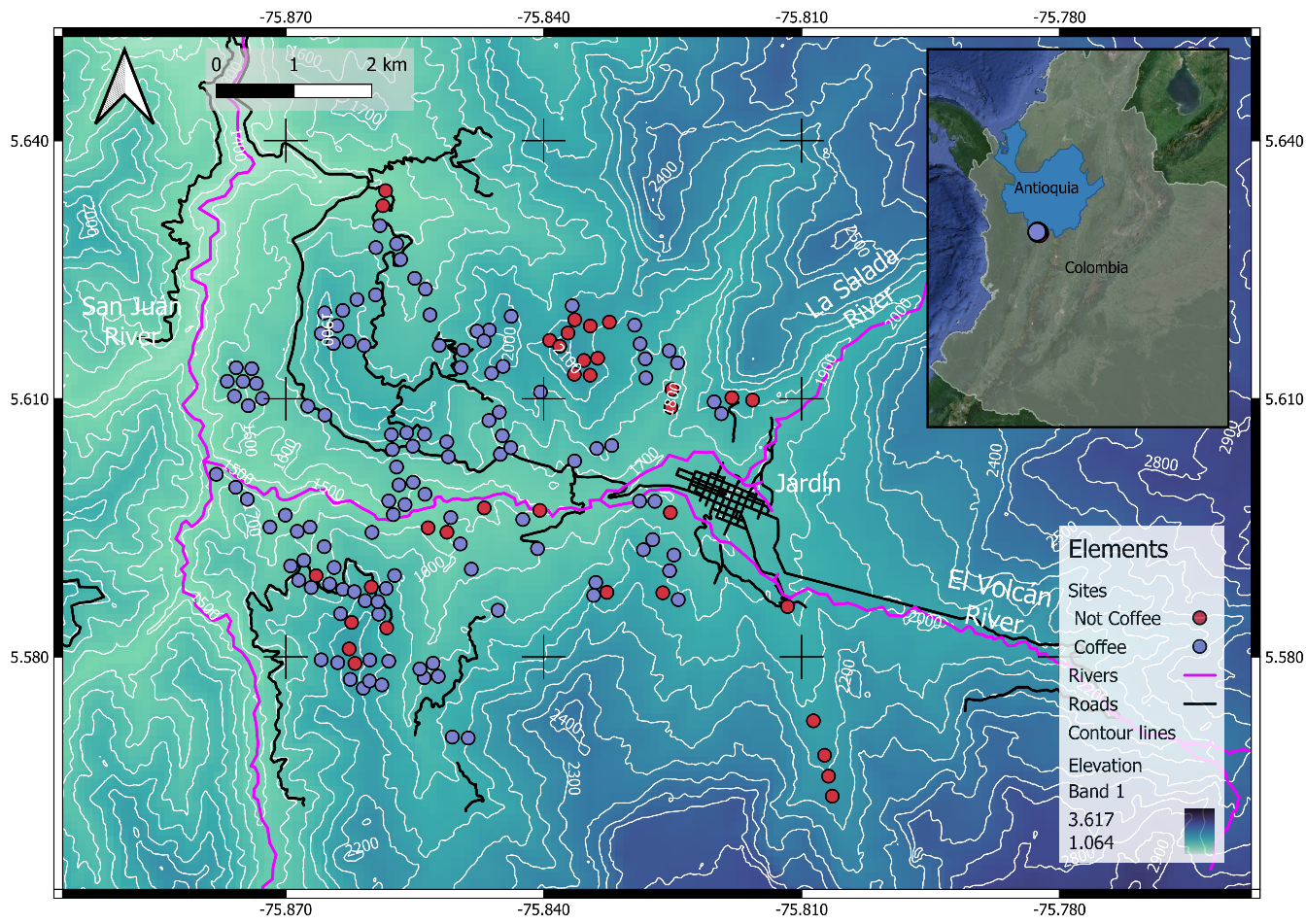
