## Supplementary material for "Differential Avian Responses to Coffee Farming Lead to Community Homogenization in a Working Landscape in Jardín, Colombia": Figure S2

Figure S2 Pearson correlation coefficients among the occupancy covariates used in the analyses. Lhfi.med.3 = average human footprint, ch.med = average canopy height, fhdpai =average foliage height diversity, elev.mean.1 = average elevation.

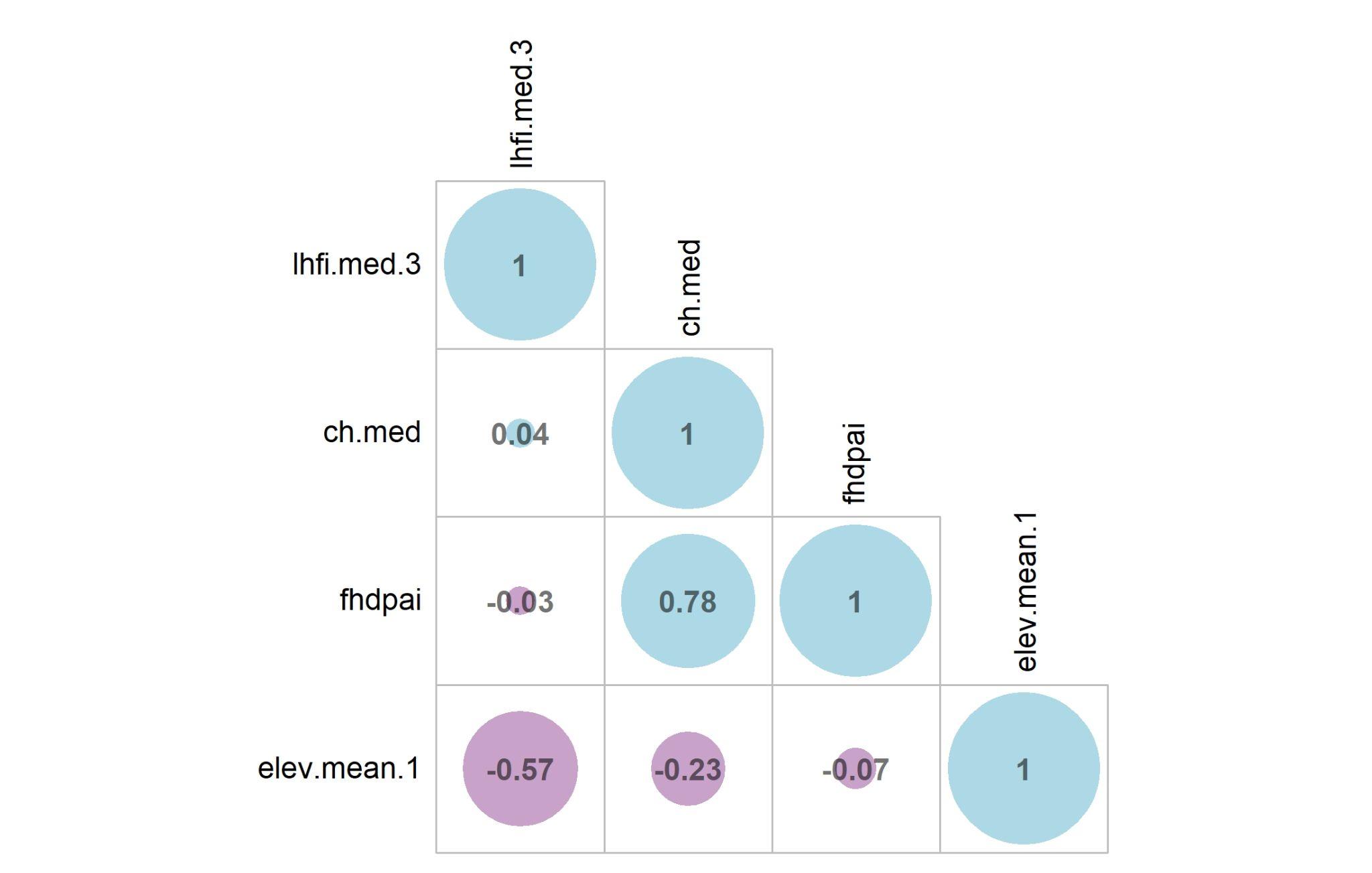
