## Supplementary material for "Differential Avian Responses to Coffee Farming Lead to Community Homogenization in a Working Landscape in Jardín, Colombia": Figure S3

Figure S3 Density distribution of the occupancy coefficients for the five covariates used in the analyses.

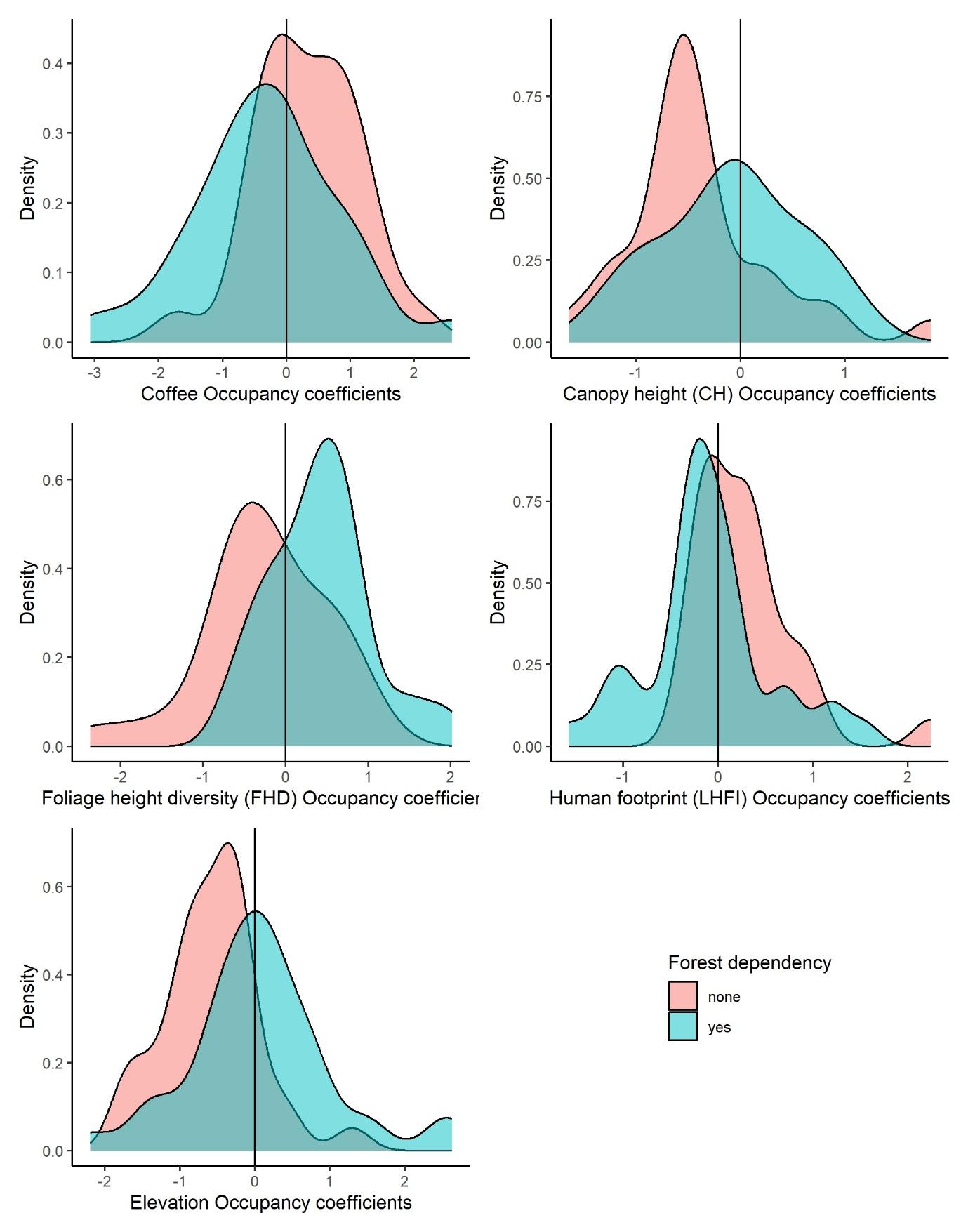
