## Supplementary material for "Differential Avian Responses to Coffee Farming Lead to Community Homogenization in a Working Landscape in Jardín, Colombia": Figure S4

Figure S4 Occupancy coefficients estimated for each species used in the analyses. 1). Foliage height diversity, 2) Canopy height, 3) Human footprint, 4) Elevation and 5) Coffee presence.

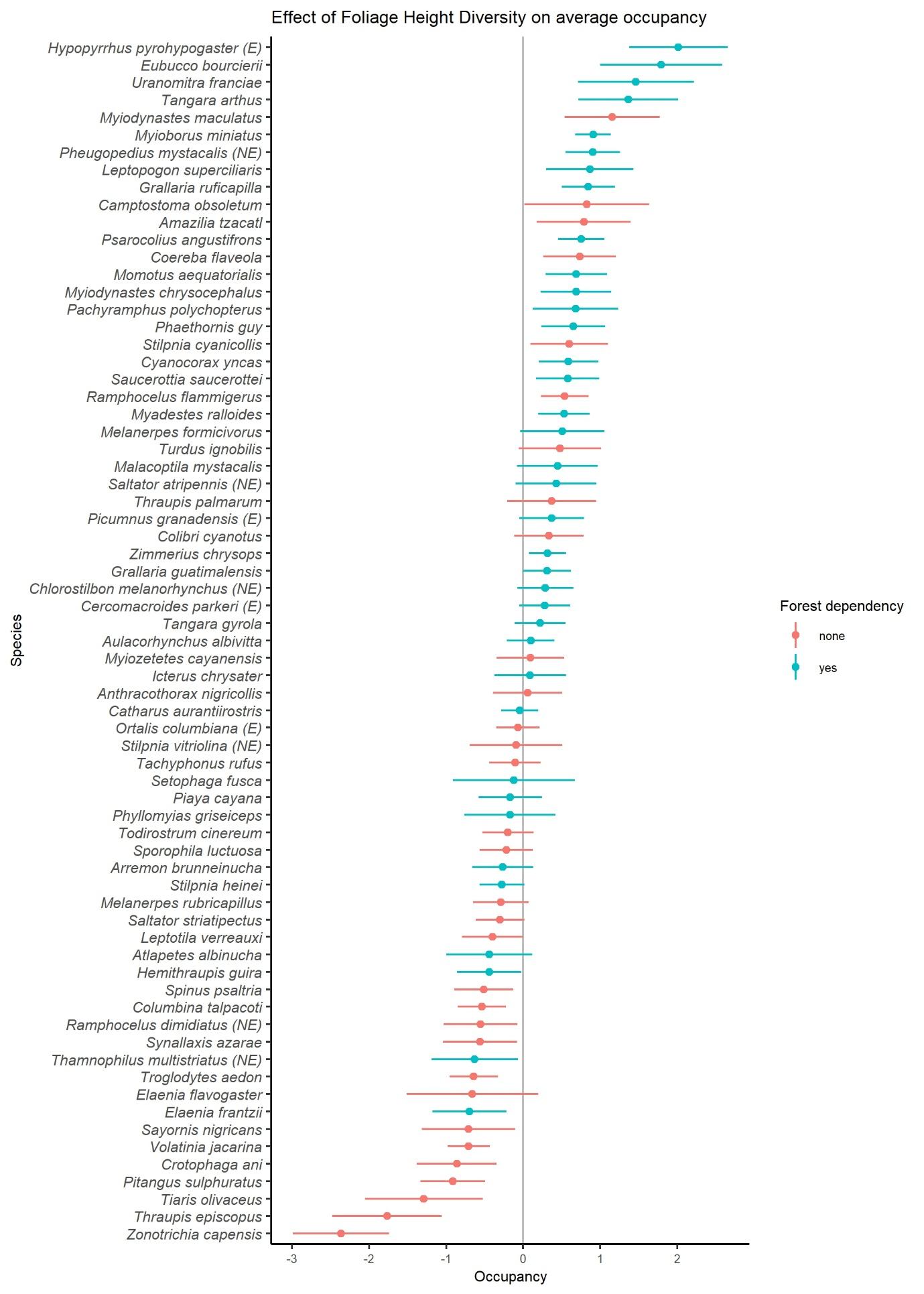

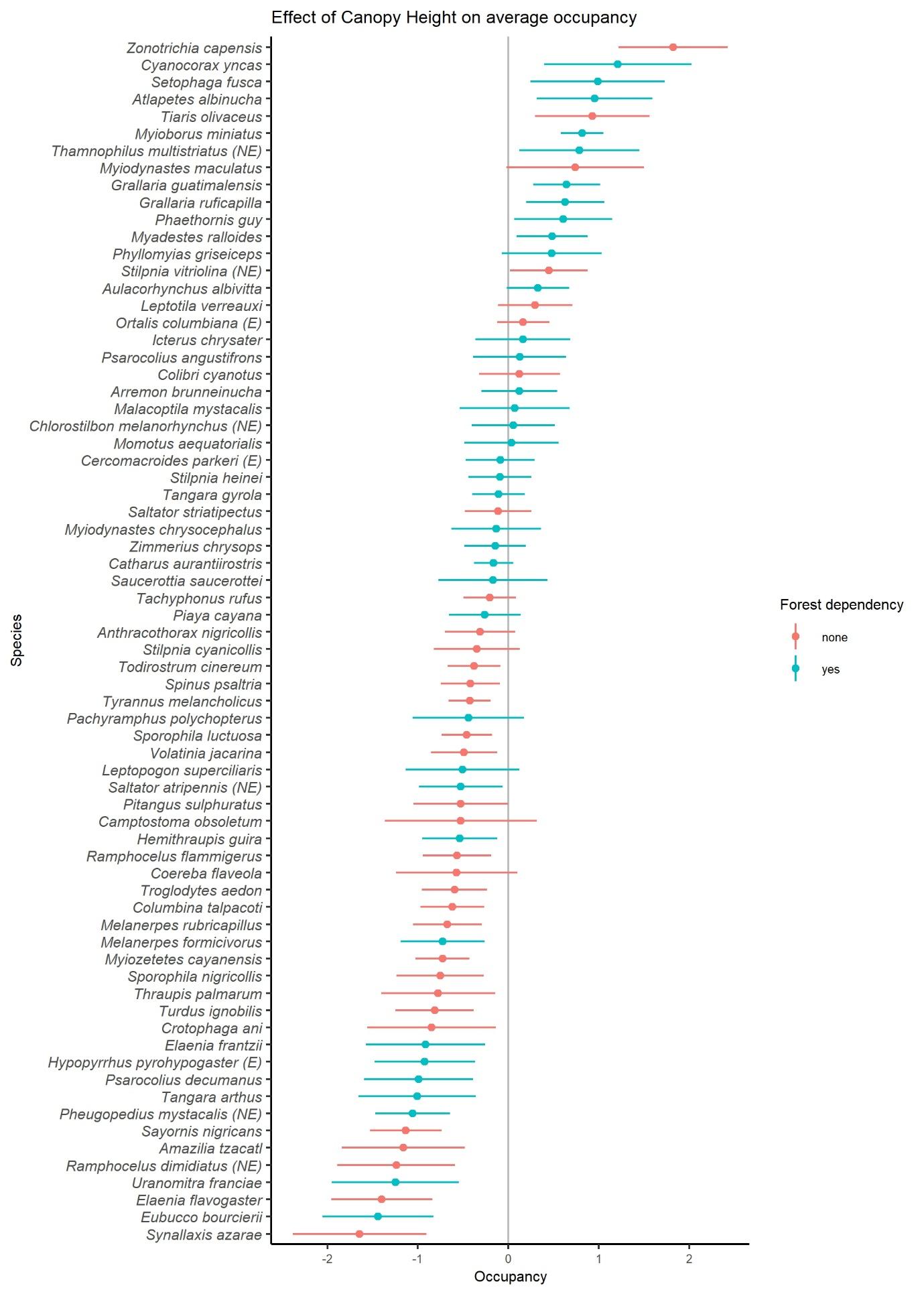

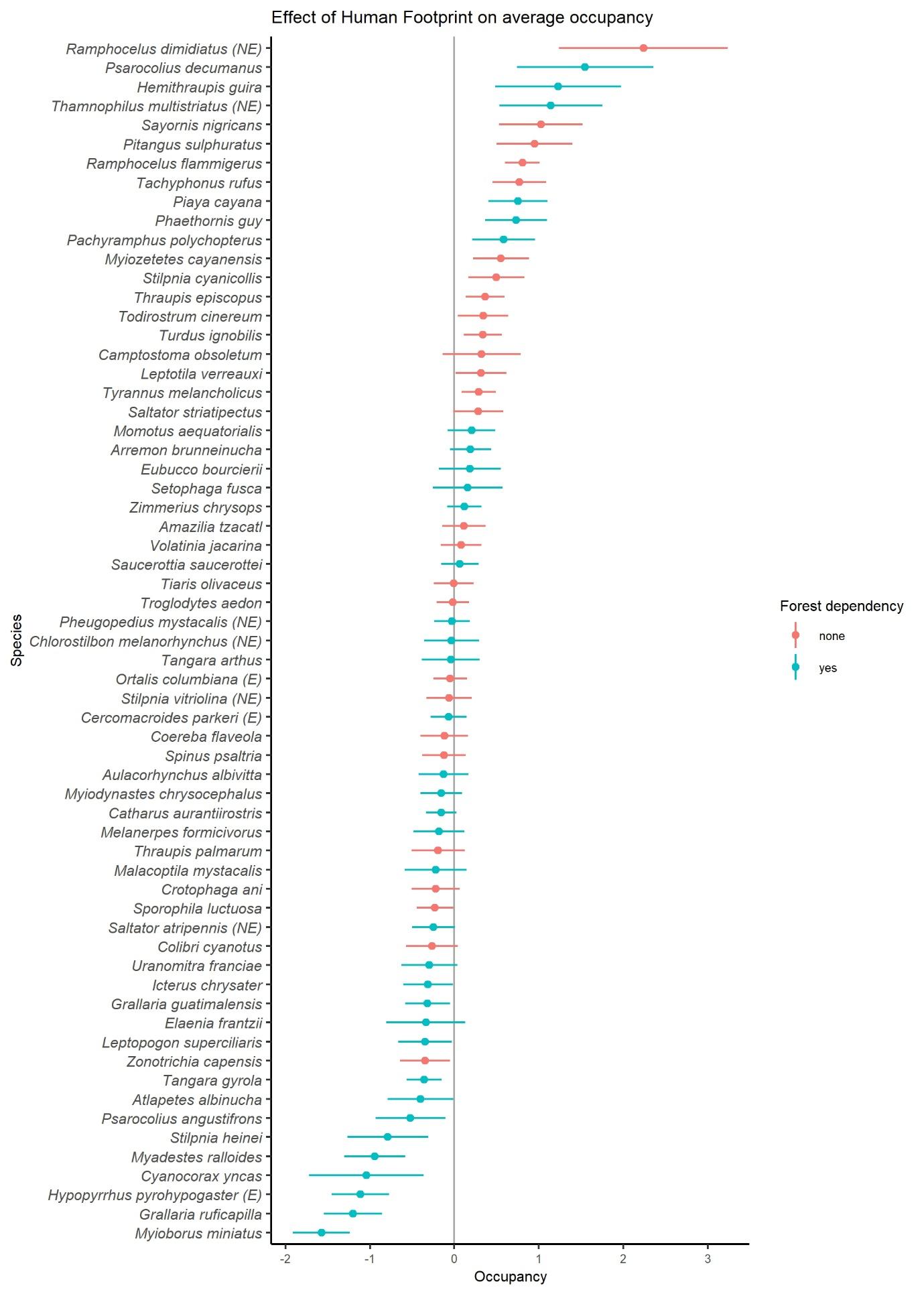

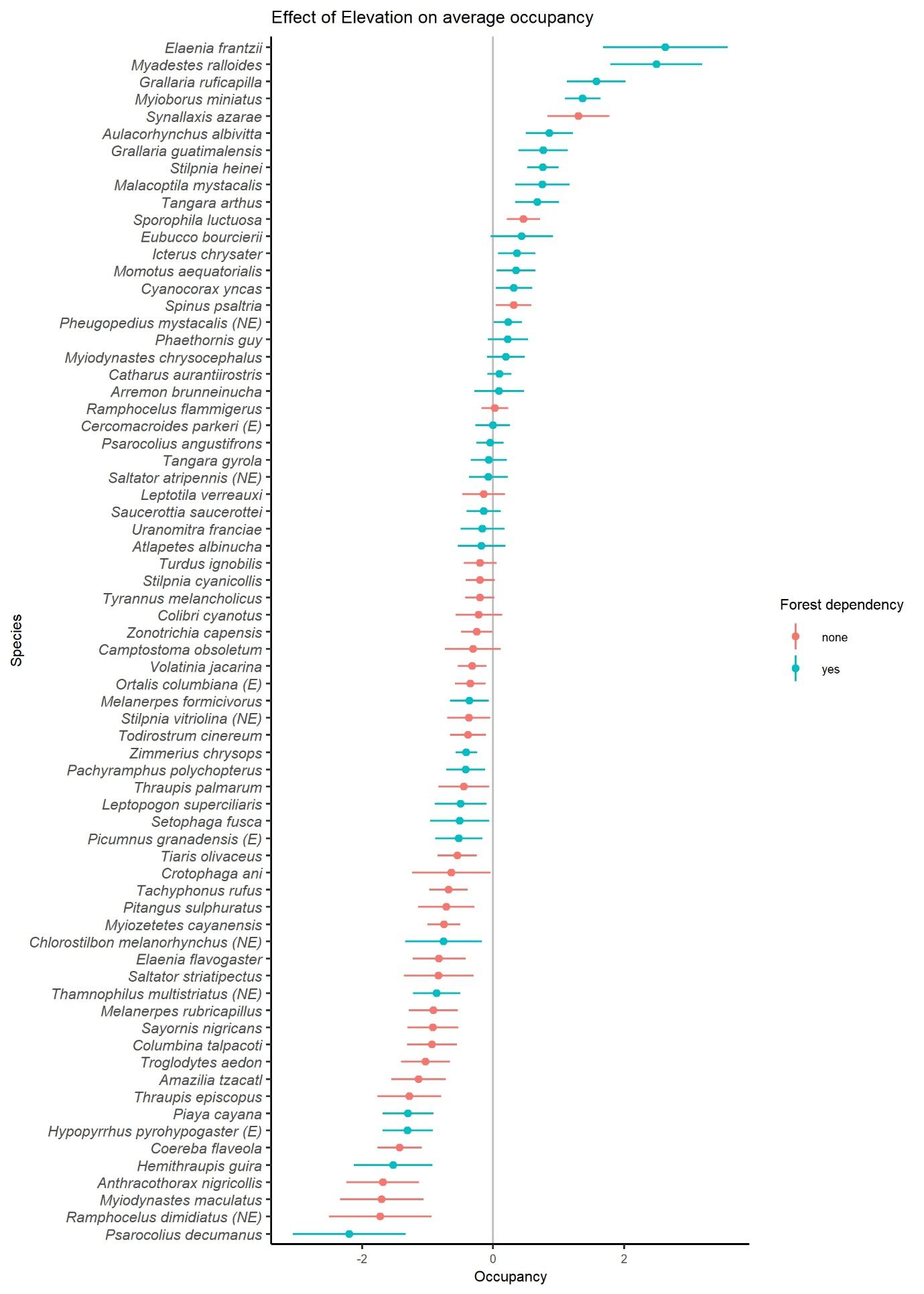

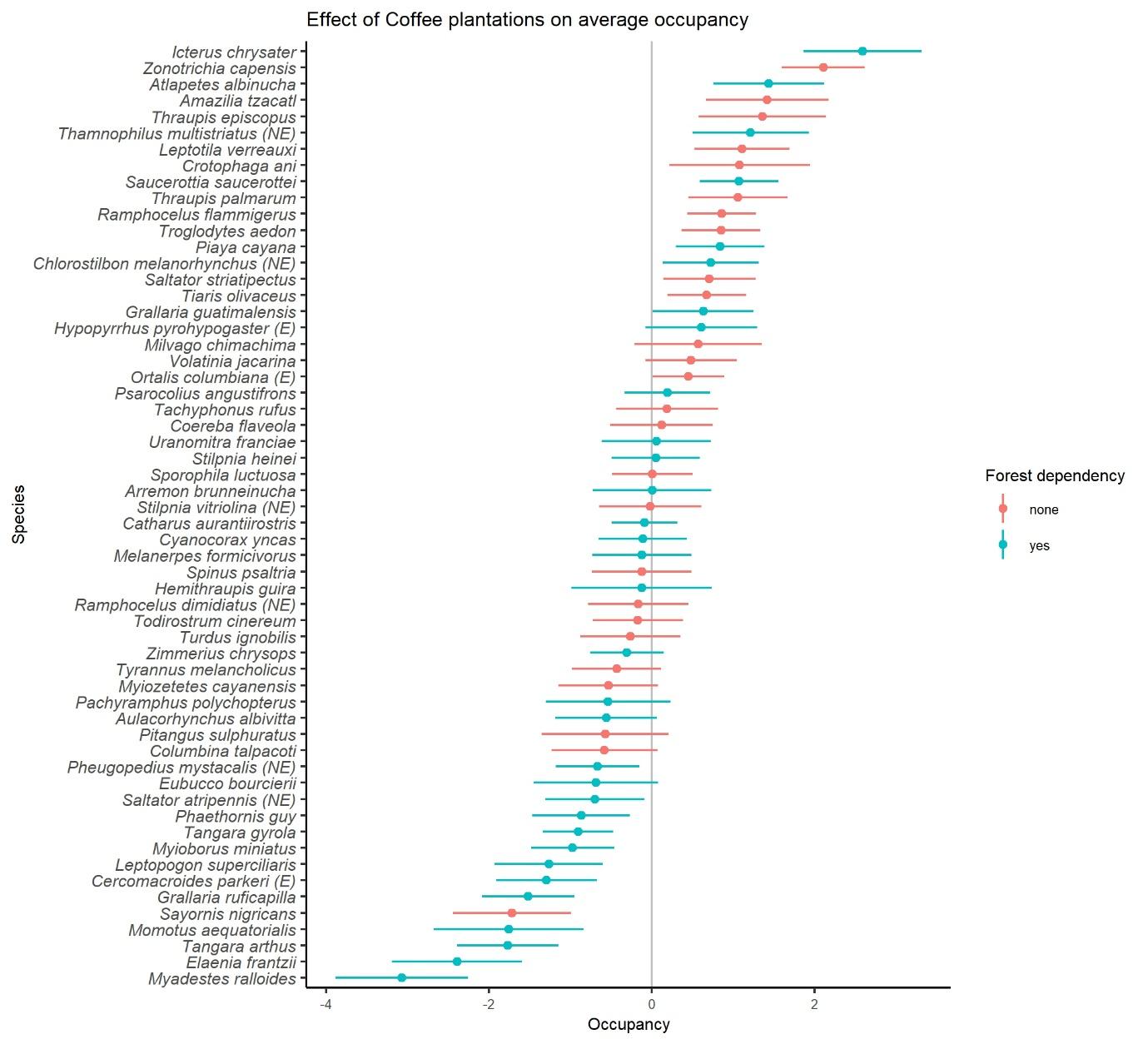
