## Supplementary material for "Differential Avian Responses to Coffee Farming Lead to Community Homogenization in a Working Landscape in Jardín, Colombia": Figure S5

Figure S5 Observed dissimilarity (Bray Curtis distance) based on the complete community of birds as a function of predicted ecological distances (including all covariates). Each dot corresponds to a site-pair comparison. Each dot is colored depending on whether it refers to a comparison between coffee sites, not-coffee sites, or a coffee site and a non-coffee site. Side graphs represent the density of points for each axis.

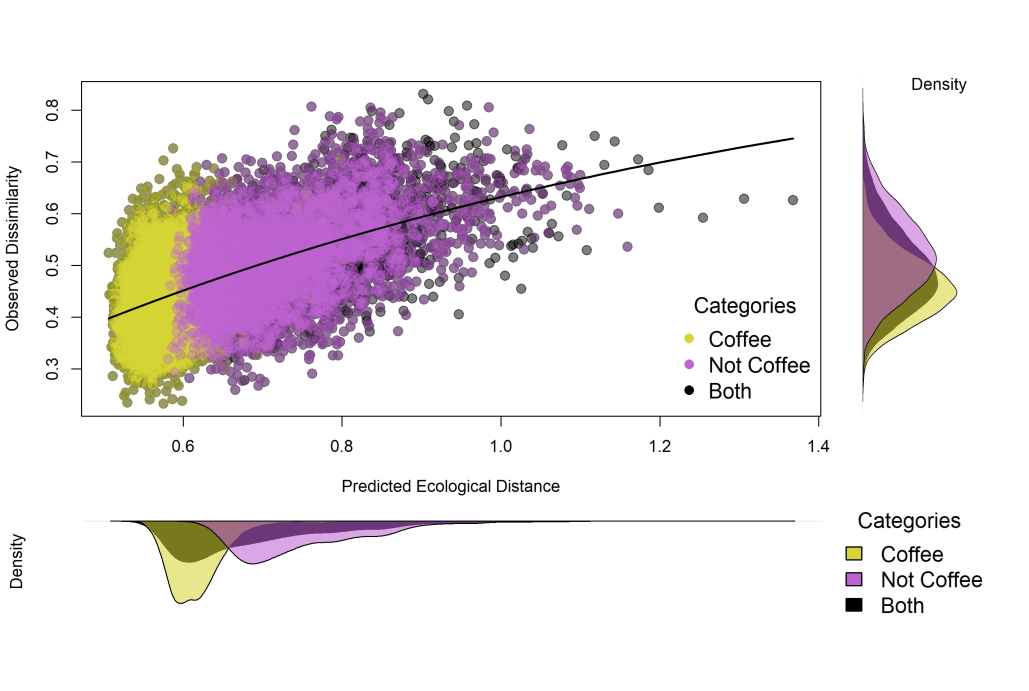
