## Supplementary material for "Differential Avian Responses to Coffee Farming Lead to Community Homogenization in a Working Landscape in Jardín, Colombia": Figure S6

Figure S6 I-splines of each covariates used in the complete community analysis. The I-splines show the relationship between each variable and the predicted ecological distance. In this case, geographic distance (Geo Dist) is the most important predictor variable. The bar-graph on the lower right is the result of the relative importance of the covariates used.

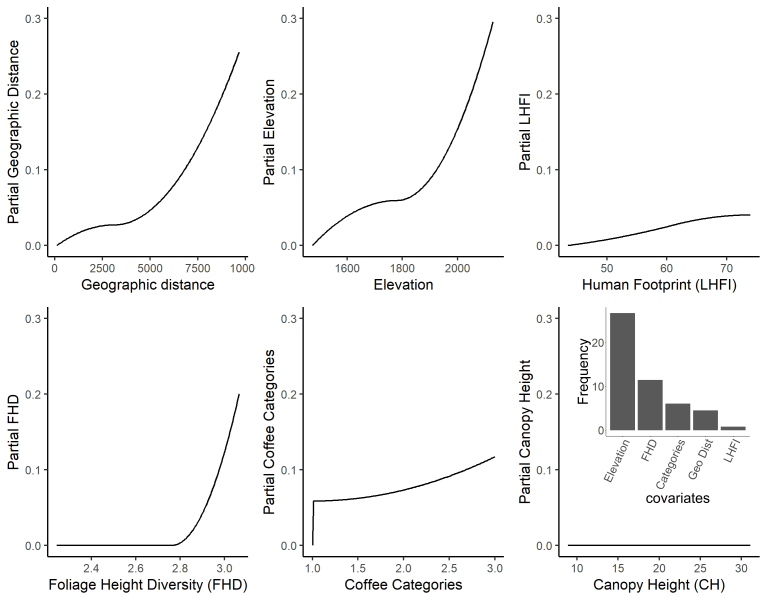
